# The microcephaly protein WDR62 regulates cellular purine metabolism through the HSP70/HSP90 chaperone machinery

**DOI:** 10.1101/2024.07.01.601630

**Authors:** Matthew J. Morris, Yvonne Y. Yeap, Chi Chen, S. Sean Millard, Julia K. Pagan, Dominic C. H. Ng

## Abstract

Inherited mutations in WD repeat-containing protein 62 (WDR62) are associated with microcephaly (*MCPH2*). While WDR62 plays important roles in mitosis and centriole biogenesis, additional WDR62 functions may cause abnormal brain growth. Here, we reveal a novel WDR62 role in the molecular chaperone network regulating purine metabolism. In response to hyperosmotic stress, WDR62 redistributes to purinosomes—phase-separated membraneless assemblies of purine metabolic enzymes and their chaperones. While WDR62 is not needed for purinosome formation, its loss disrupts purine homeostasis, resulting in the accumulation of purine nucleotide intermediates and a reduction in the levels of hypoxanthine-guanine phosphoribosyl transferase (HPRT), a key purine salvage enzyme. We link this to WDR62’s interaction with Bcl2-associated athanogene 2 (BAG2), a co-chaperone that modulates the function of HSP70/90. In cells lacking WDR62, BAG2 levels are elevated and HPRT stability is reduced. Knocking down BAG2 in these cells restores HPRT levels, underscoring the crucial role of WDR62-BAG2 interactions in chaperone-mediated stability and turnover of metabolic pathway enzymes. Notably, common microcephaly-associated mutations in WDR62 alter its interaction with BAG2, suggesting that purine metabolic defects resulting from WDR62 mutations may underlie microcephaly in humans.

## Introduction

Microcephaly is a complex neurological condition attributed to inherited mutations or prenatal exposure to infection, metabolic imbalances, or chemical toxicity [1]. One commonly mutated microcephaly gene, *MCPH2,* encodes WD40 repeat-containing protein 62 (WDR62), a widely expressed protein with vital non-redundant functions in embryonic brain growth [2–4].

WDR62 exhibits a dynamic subcellular distribution that is coordinated with the cell cycle [5, 6]. WDR62 was initially characterised as a c-Jun N-terminal kinase (JNK)-binding protein involved in stress-activated signal transduction [7]. During mitosis, WDR62 is associated with spindle microtubules and is required for spindle organization, mitotic progression, and division polarity [5, 8–12]. The ability of WDR62 to regulate the orientation of mitotic divisions may have significant consequences on daughter cell fate, particularly during the proliferation of progenitor cell populations [11, 13]. Additionally, a proportion of WDR62 localises to centrosomes, where it contributes to centriole biogenesis and ciliogenesis [9, 14]. Consequently, loss or mutation of WDR62 results in abnormal centrosome numbers that may cause genetic instability or alter primary cilia dynamics—both of which define the self-renewal capacity of progenitor cells [9, 15]. Thus, WDR62 is a multi-faceted protein crucial for cell survival, proliferation, differentiation, and the integration of environmental cues.

Despite the complex functions attributed to WDR62 since it was first identified, it remains unclear precisely how WDR62 mutations lead to microcephaly phenotypes. For example, WDR62-related spindle orientation defects may not be sufficient to induce defects in mitotic index, cell cycle delay, and cortical neurogenesis [14, 16–20]. Further, we previously reported that neuroblast-specific depletion of WDR62 in *Drosophila* did not substantially change brain volume, while the loss of WDR62 in the glial lineage reduced brain volume without observable effects on spindle orientation [21]. Similarly, misoriented divisions in neural stem cells due to the loss of polarity determinant aPKCλ or spindle pole protein LGN (Gpsm2) do not consistently alter brain volume in mice [22, 23]. Given that cytoskeletal, centrosomal, and cell cycle processes are interconnected, determining whether WDR62 mutant phenotypes arise from direct or indirect effects is challenging.

A large proportion of WDR62 is cytosolic during interphase, rather than being associated with microtubules and centrosomes [4]. Despite well-described functions for microtubule and centrosome associated WDR62, the role of cytosolic WDR62 has remained largely uncharacterized. Using BioID to identify novel interactors, we found that the WDR62 interactome comprised molecular chaperones such as HSP70, HSP90, and their co-regulators, BAG2, STIP1 and DNAJC7. This chaperone network regulates the stability, turnover and higher-order assembly of a diverse range of client proteins, including enzymes involved in purine metabolism [24–26].

Here, we demonstrate that WDR62 and BAG2 interactions are altered by microcephaly associated WDR62 mutations. In response to cell stress, WDR62 was recruited to phase-separated *purinosomes*, that comprise *de novo* purine synthesis enzymes and molecular chaperones, and not to stress granules that contain halted translational machinery. WDR62 loss caused accelerated degradation of a critical enzyme involved in purine synthesis, hypoxanthine-guanine phosphoribosyl transferase (HPRT), and led to the accumulation of purine nucleotide intermediates. We linked these defects to dysregulated chaperone activity (HSP90 and BAG2) required for HPRT stability. Our findings unveil a novel role for WDR62 in maintaining the stability and activity of purine metabolic enzymes and overall purine homeostasis, which is implicated in neuronal differentiation, migration, and neurodevelopment [27–30]. We propose that this role in purine homeostasis may further explain how mutations in WDR62 lead to microcephaly.

## Results

### WDR62 interactors include a network of molecular chaperones and their co-regulators involved in protein folding

To identify novel WDR62 binding partners, we generated WDR62 fusion proteins tagged with the BirA* promiscuous biotin ligase. This allowed for spatially restricted biotin labelling, affinity isolation, and detection of proximal proteins and transient interactors within an approximate 10 nm radius (BioID) of WDR62 (Fig. 1A) [31]. Exogenously expressed WDR62-BirA*-HA and WDR62-HA were localized to the cytosol and spindle poles in interphase and mitotic cells, respectively (Fig. 1B), resembling the spatial distribution of endogenous WDR62 [4]. We validated, using fluorophore-conjugated streptavidin, that biotin labelling occurred only in cells expressing BirA* fusion proteins and following biotin feeding (Fig. 1B). We also confirmed that WDR62-BirA*-HA was localised in proximity to established WDR62 binding partners such as JNK1/2, CEP170, AURKA, and MAPKBP1 using proximity-labelling followed by immunoblotting (Fig. 1C).

**Figure 1.**
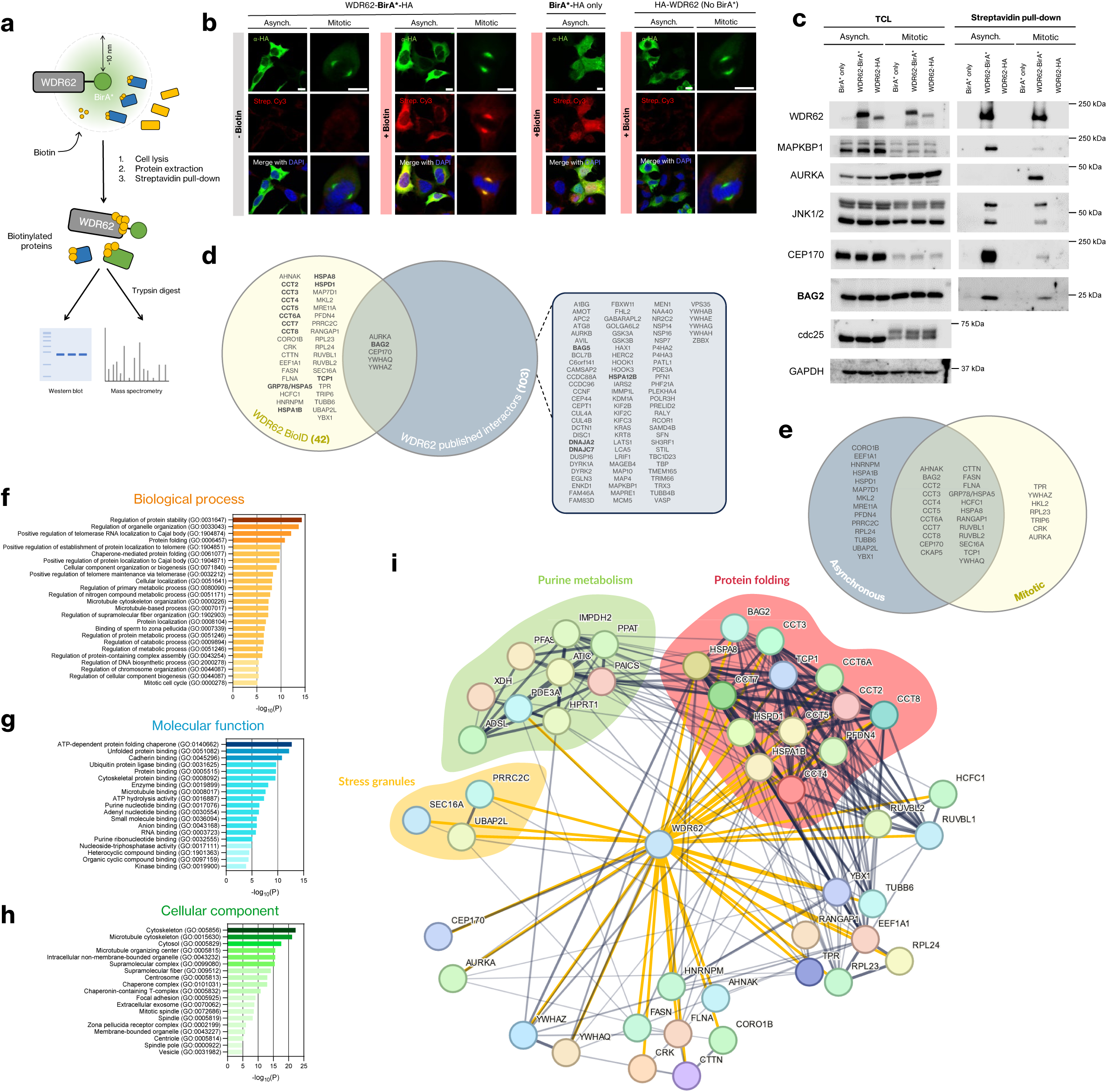
BioID proximity-interaction analysis of WDR62. **(A)** Schematic of the BioID study of WDR62-interacting proteins. **(B)** Validation of WDR62-BirA*-HA localisation in AD293 cells by fluorescence microscopy. **(C)** Streptavidin pull-down and immunoblot confirms MAPKBP1, AURKA, JNK1/2, CEP170 and BAG2 as interactors of WDR62 in asynchronous and mitotically arrested cells. **(D)** Venn diagram of published WDR62 interactors (BioGRID database), interactors identified through BioID screening, and interactors found in both datasets. **(E)** BioID-identified interactors sorted by those identified in asynchronous or mitotic cells, or both. **(F-H)** Gene ontology (GO) analysis of significantly enriched **(F)** biological processes, **(G)** molecular functions, and **(H)** cellular components (-log_10_(P)) mediated/localised by WDR62 interactors. **(I)** STRING-based reconstruction of the network of proteins identified as interactors of WDR62 by our BioID study. Yellow lines indicate empirically confirmed interactions, grey lines are interactions curated in the STRING protein interaction database.

Satisfied that WDR62-BirA*-HA behaved equivalently to endogenous WDR62, we performed BioID proximity-labelling screens in asynchronous or mitotically arrested (low dose nocodazole) cells, to distinguish cell-cycle dependent proximity partners, isolated biotinylated proteins by streptavidin pulldown, and identified proteins in the pulldown fraction by LC-MS/MS. Overall, we identified 42 interactors, with 8 of these matching entries in the BioGRID database, which compiled 103 interactors from a range of studies using affinity capture, yeast two-hybrid and other biochemical analyses of protein-protein interactions (Fig. 1D). Mitotic arrest of cells led to the identification of a limited set of additional interactors including AURKA (Fig. 1E). Gene ontology (GO) analyses indicated that WDR62 proximal partners were significantly enriched for proteins localized to the cytosol and microtubule cytoskeleton and involved in the regulation of protein stability and homeostasis, with an emphasis on chaperone-mediated protein folding (Fig. 1F-H). Of note, proximity labelling identified members within the chaperonin-containing TCP1 complex (CCT2-CCT8, TCP1), the stress-responsive molecular chaperones HSP60 and HSP70, as well as several HSP40 family members (e.g., BAG2, BAG5, DNAJA2, DNAJC7) which function as HSC70/90 and HSP70/90 co-chaperones under basal and stress-activated conditions (Fig. 1I) [32]. Interestingly, WDR62-BirA* labelling of BAG2 was reduced in mitotically arrested cells compared to asynchronous populations (Fig. 1C). As BAG2 expression levels were unchanged in mitosis, unlike CEP170 and AURKA, we conclude that WDR62 interactions with BAG2 were reduced in mitotic cells and increased during interphase (Fig. 1C). Together, these results demonstrate that WDR62 interacts with protein chaperone complexes regulating protein folding and stability.

### WDR62 microcephaly mutations alter binding to BAG2

Having identified BAG2 in both our BioID study and among the published interactors of WDR62, we investigated its interaction with WDR62 in more detail using AlphaFold2-Multimer (AF2) to predict heterodimeric WDR62-BAG2 structures. Considering that WDR62 contains long disordered domains, our initial approach involved evaluating the validity of predicted structures through predicted alignment error (PAE) plots. Our primary objective was to identify residues with low error scores (high confidence), particularly those involved in a predicted protein-protein interfaces. The predicted structural model of WDR62 showed that its WD repeat domains fold into two adjacent seven-bladed beta-propeller structures, predicted with high confidence (mean pLDDT: 76.34) (Fig. 2A, B). Conversely, most of the C-terminal half of WDR62 exhibited low confidence scores (mean pLDDT: 35.74), indicating disorder and the absence of a fixed three-dimensional structure (Fig. 2A, B). The predicted WDR62-BAG2 heterodimer demonstrated that BAG2 interacts with one of the two beta-propeller domains of WDR62, which contain WD repeat motifs (Fig. 2A). Additionally, in most predicted models, the C-terminal tail of WDR62, which includes the α3 helix-loop-helix domain [33], appeared to interact with BAG2 on the side opposite its interface with the WD repeat domains (Fig. 2A). Indeed, heterodimeric structure predictions of BAG2 with a microcephaly-identified mutant of WDR62 lacking this helix-loop-helix domain (WDR62(3936dupC), aa. 1-1329), demonstrated a reduction in interphase area (Å^2^) in comparison to full-length WDR62 (Fig. S1). This observation suggests that BAG2 binds WDR62 primarily through its WD repeat domains and may rely on the C-terminal helix-loop-helix domain for additional stability.

**Figure 2.**
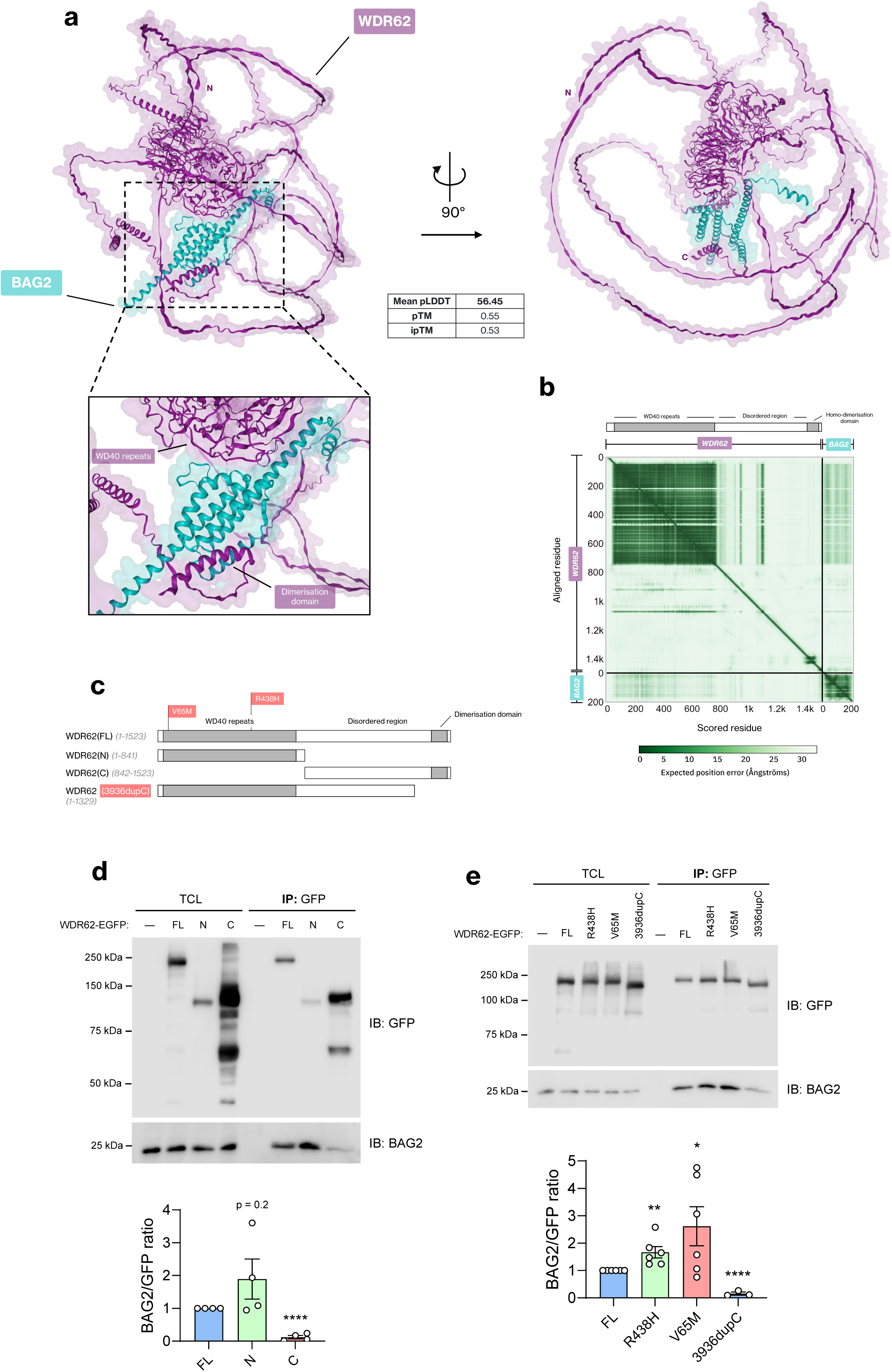
Microcephaly-associated mutations in WDR62 alter binding to BAG2. **(A)** Top-ranked model predicted by AlphaFold2 between WDR62 and BAG2 with calculated mean predicted local distance difference test (pLDDT), predicted template modelling (pTM) and interphase predicted template modelling (ipTM) scores. **(B)** Predicted aligned error (PAE) plot of predicted WDR62-BAG2 heterodimer. X- and Y-axes show indexed residues of corresponding subunits, as indicated. Aligned error in angstroms (Å) is colour coded (green = low PAE (high confidence), white = high PAE (low confidence). **(C)** Schematic depicting WDR62(FL) (aa. 1-1523), truncated mutants WDR62(N) (1-841), WDR62(C) (842-1523), and microcephaly-associated mutants WDR62(3936dupC) (1-1329), V65M and R438H. **(D)** GFP-trap immunoprecipitation of WDR62(FL)-, WDR62(N)- and WDR62(C)-EGFP and immunoblot for GFP and endogenous BAG2. **(E)** GFP-trap immunoprecipitation of WDR62(FL)-, WDR62(R438H)-, WDR62(V65M), and WDR62(3936dupC)-EGFP and immunoblot for GFP and endogenous BAG2. Densitometric quantification of BAG2/GFP ratio for **(D)** and **(E)** represented underneath blots. Data represent mean ± SEM of *n* > 3 independent replicates.

We next sought to empirically validate these predictions and determine which WDR62 regions were required for binding BAG2. To this end, we assessed the interaction of BAG2 with EGFP-tagged full-length WDR62 (WDR62(FL)-EGFP), in addition to N-terminal (WDR62(N)-EGFP, aa. 1-841) and C-terminal fragments (WDR62(C)-EGFP, aa. 842-1523) corresponding to the WD40 repeat and disordered regions of WDR62, respectively (Fig. 2C). We isolated EGFP-tagged proteins with GFP-trap agarose and assessed co-precipitation of endogenous BAG2 by immunoblot. Consistent with our findings from the BioID labelling system and streptavidin pulldown, we observed that endogenous BAG2 interacts with full-length WDR62 (Fig. 2D). In addition, consistent with our AF2 prediction, BAG2 co-immunoprecipitated with WDR62(N)- EGFP but not WDR62(C)-EGFP (Fig. 2D), suggesting an interaction with the N-terminal half of WDR62 enriched in WD repeat motifs. Given that chaperones typically bind to unfolded regions of clients [34], the interaction of BAG2 with the ordered WD repeat regions of WDR62 suggest that WDR62 is likely a binding partner of BAG2 and not a client or substrate.

We next determined whether patient-identified mutations in WDR62 affected BAG2 binding. These included two missense mutants (R438H and V65M) [9, 11], and the frameshift mutant (3936dupC) [3, 4, 35], mentioned above (Fig. 2C). Interestingly, we found that the R438H and V65M mutants immunoprecipitated significantly more BAG2 compared to wild-type WDR62 (Fig. 2E). Thus, the N-terminal region of WDR62 is sufficient for binding BAG2 and this is altered by microcephaly-associated missense mutations. In contrast, the 3936dupC frameshift mutant of WDR62, led to a substantial decrease in BAG2 interaction, consistent with our AF2 prediction (Fig. 2E). This indicates that, while the disordered C-terminal half of WDR62 is not sufficient to bind BAG2, the C-terminal tail comprising the helix-loop-helix domain—required for WDR62 dimerisation—is required for BAG2 to bind full-length WDR62. These results indicate that patient-identified mutations in WDR62 substantially alter interactions with BAG2, which suggest WDR62 roles within the chaperone network may intersect with brain development.

### WDR62 assembles granules that are distinct to conventional stress granules

BAG2 has established functions as a HSP70 co-chaperone with nucleotide exchange factor (NEF) activity that accelerates the protein folding cycle [36, 37]. In addition, it was recently revealed that the phase-separation of BAG2 into stress-induced condensates directed client proteins for ubiquitin-independent proteasomal degradation in response to cell stress [38]. Similarly, WDR62 was previously reported to be recruited to cytoplasmic stress granules which are also formed through phase separation [7, 39].

To interrogate the spatial distribution of WDR62 following stress stimulation, we transiently expressed WDR62-mCherry in AD293 cells. Under basal conditions, WDR62-mCherry was uniformly distributed throughout the cytoplasm (Fig. 3A). Upon acute exposure to hyperosmotic stressors, including sugars such as dextrose or sucrose, or polyols such as sorbitol (0.5 M, 1 h), WDR62 assembled distinct cytoplasmic puncta or granules (Fig. 3A). As a control, we confirmed that mCherry alone, or the WDR62 paralog MAPKBP1 do not form puncta under identical stress conditions (Fig. S2, S3). We observed the formation of WDR62 granules at sorbitol concentrations exceeding 50 mM, with a gradual increase in WDR62 condensation as we incrementally raised the concentration over a 10 min period (Fig. 3B). WDR62 granules exhibited homogenous staining throughout their z-volume as shown by three-dimensional orthogonal projections (Fig. 3C). WDR62 granules ranged from spherical to irregular in shape with an average circularity of 0.58 ± 0.02 (Fig. 3D). In response to sorbitol treatment, the majority (>90%) of cells assembled WDR62 granules (Fig. 3E), with an average number of 102.8 ± 10.2 granules per cell that were on average 0.86 ± 0.15 µm in diameter (Fig. 3F).

**Figure 3.**
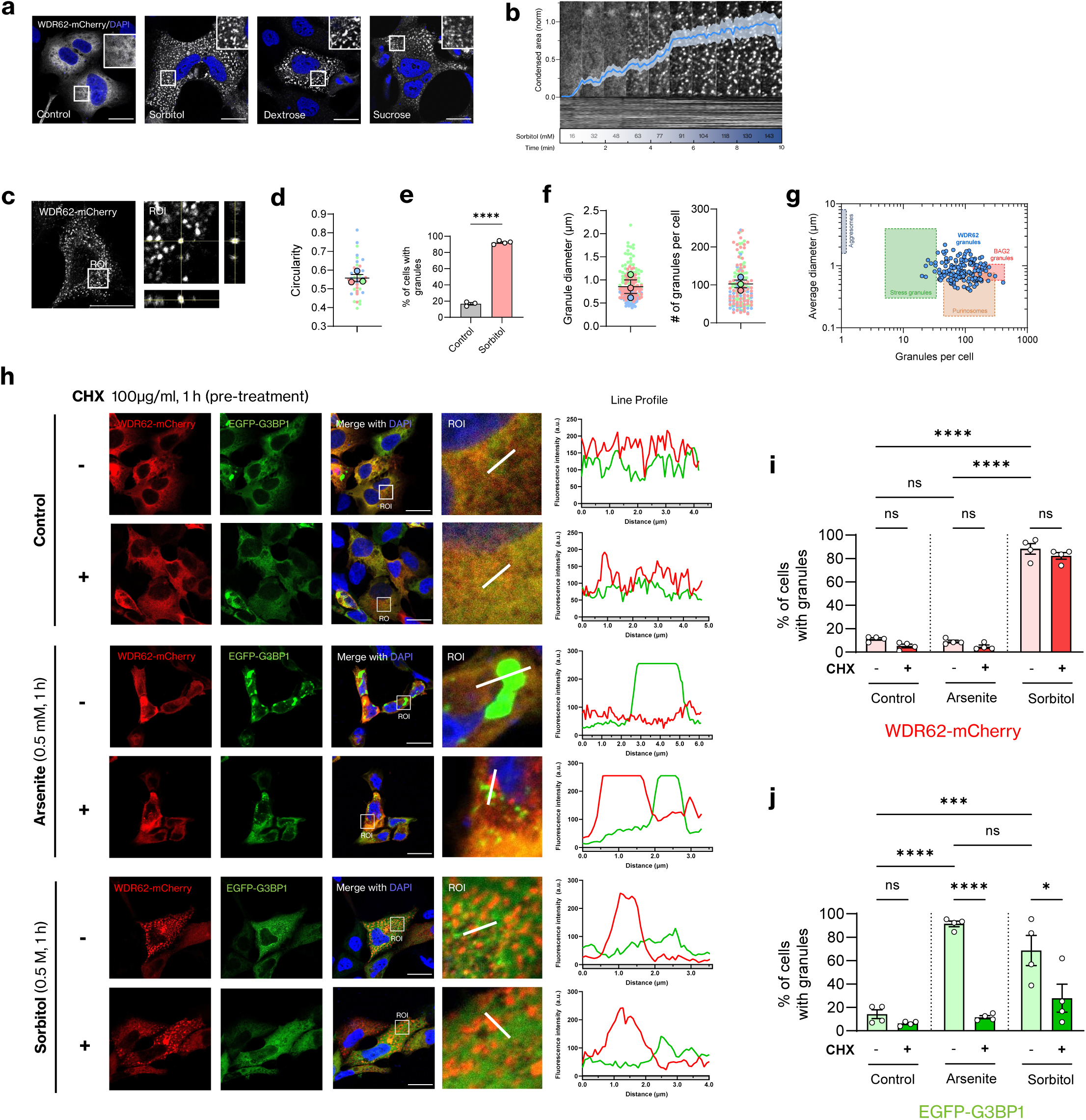
WDR62 assembles into stress granule-independent condensates following hyperosmotic stress. **(A)** Representative images of AD293 cells expressing WDR62-mCherry, showing WDR62 de-mixes into cytoplasmic puncta following treatment with 0.5 M sorbitol, dextrose, or sucrose for 1 h. **(B)** Titration of WDR62 granule assembly. A concentrated solution of sorbitol was added dropwise to a live-cell culture dish containing AD293 cells expressing WDR62-mCherry. Sorbitol was added every minute for the duration of the experiment (10 min). Total sorbitol concentration in culture media is displayed in the figure. Blue line represents a quantification of the total WDR62 condensation area over time. Condensed area is the total sum of WDR62-mCherry expression normalised by its highest value. Scale bar represents 5 µm. **(C)** 3-dimensional orthogonal projection of WDR62 granules. **(D)** Average circularity of WDR62 granules. A circularity of 1 is equivalent to a perfect circle. **(E)** The average proportion (%) of cells containing WDR62 granules before and after sorbitol stress. Two-tailed unpaired Student’s T-test (****p<0.0001). **(F)** The average diameter (µm) and number per cell of WDR62 granules. **(G)** XY plot depicting average diameter and granules per cell of WDR62 granules (n=180 cells) compared to reported sizes of aggresomes, stress granules, purinosomes and BAG2 granules. **(H)** AD293 cells transiently transfected with WDR62-mCherry and stress granule marker, EGFP-G3BP, pre-treated with 100 µg/ml cycloheximide (CHX) (+) or vehicle control (-) for 1 h. Cells subsequently treated with 0.5 mM sodium arsenite or 0.5 M sorbitol. Representative confocal micrographs demonstrate minimal co-localisation between WDR62-mCherry and G3BP-EGFP, indicated by fluorescence intensity plots (y-axis represents fluorescence intensity (a.u.), x-axis (µm) represents length of white line drawn on ROI image. **(I)** Bar graph representing proportion (%) of cells containing WDR62 granules following treatments in (A). **(J)** Bar graph representing proportion (%) of cells containing WDR62 granules following treatments in (A). One-way ANOVA with Tukey’s multiple comparisons. (*p<0.05, ***p<0.001, ****p<0.0001, n.s. is p>0.05). All scale bars represent 20 µm. *n* = 4 biological replicates.

To determine if the effect of sorbitol on WDR62 was due to osmotically-induced cell volume reduction, we pre-treated cells with the osmoprotectant trimethylglycine (betaine, 20 mM) to stabilise cell volume [40, 41], and showed that this significantly reduced WDR62 granule formation compared to cells treated with sorbitol alone (Fig. S4A, B). However, we found that in response to hyperosmotic media containing the ionic osmolyte sodium chloride (NaCl) at 0.25 M (500 mOsm), cells robustly assembled G3BP-positive stress granules but the distribution of WDR62 remained diffuse (Fig. S4C). This suggests that WDR62 assembles granules in response to non-ionic osmolytes, as a function of osmolarity and not an increase in salt concentration.

Our findings with NaCl, inducing similar osmotic pressure, also indicated that cell volume changes alone were not sufficient to induce formation of WDR62 granules. Although cell membrane permeability to polyols is low, sorbitol permeases have been identified in human erythrocytes and renal epithelial cells, allowing intracellular sorbitol uptake [42, 43]. Indeed, sorbitol, and other sugars such as trehalose and sucrose, may alter protein-solvent interactions and modulate protein stability *in vitro* [44, 45]. To determine if bioactive properties of sorbitol contributed to WDR62 granule assembly, we treated cells with millimolar concentrations of sorbitol (5 or 50 mM, 1 h), either with or without 0.25 M NaCl. In this context, the addition of sorbitol in millimolar concentrations only minimally increased osmolarity (+5 or +50 mOsm), compared to 0.25 M NaCl (+500 mOsm). While NaCl alone does not affect the cytoplasmic distribution of WDR62 (Fig. S4A, F), the addition of low-dose sorbitol in combination induced a significant increase in granule assembly (Fig. S4D-F). We conclude that non-ionic osmolytes such as sorbitol promote the assembly of WDR62 granules due to cell volume changes combined with an undetermined bioactive effect specific to these osmolytes (Fig. S4G).

When compared to other cytoplasmic granules [38, 46–48], WDR62 granules exhibit similar diameters and per cell numbers to BAG2 condensates and purinosomes but less so to G3BP1-positive stress granules (Fig. 3G). We also observed that like purinosomes, WDR62 granules exhibited close association with mitochondria and microtubules [49] (Fig. S5). To determine if sorbitol-induced WDR62 granules represented conventional SGs, which are characteristically nucleated following the release of messenger ribonucleoproteins (mRNPs) from polysomes, we assessed WDR62 granule formation following treatment with the translation elongation inhibitor, cycloheximide (CHX), which is known to trap mRNPs in polysomes and selectively inhibit SG formation [50]. In AD293 cells co-expressing WDR62-mCherry and EGFP-G3BP1, arsenite treatment induced formation of EGFP-G3BP1 granules but had minimal effect on WDR62 (Fig. 3H-J). In addition, we observed minimal overlap between WDR62 and G3BP1 in the context of either sorbitol or arsenite treatment (Fig. 3H), consistent with previous reports [7]. Importantly, while CHX treatment abolished G3BP1-positive SG formation induced by arsenite and sorbitol (Fig. 3J), CHX did not prevent sorbitol-induced WDR62 granule assembly (Fig. 3I). This indicates that WDR62 granules are unlike conventional SGs as they do not require release of RNPs from stalled polysomes and do not overlap with SG markers such as G3BP1.

### WDR62 granules are dynamic and liquid-like

Stress-responsive cytoplasmic granules represent membraneless, liquid-like, biomolecular condensates assembled through liquid-liquid phase separation of protein and/or nucleic acid components [51]. To determine if WDR62 granules displayed liquid-like properties typical of biomolecular condensates, we first examined the dynamics of WDR62 granule assembly and disassembly. Through live-cell confocal imaging, we determined a robust and highly rapid diffuse-to-punctate redistribution of WDR62 within 10 s of sorbitol exposure (Fig. 4A). Further, we also observed that WDR62 granules undergo fission and fusion events (Fig. 4B). Fig. 4B-D shows one such event, where granule 1a dissociates from granule 1 and fuses with granule 2 (Fig. 4C). The area of granule 1 decreases, and granule 2 increases, over the duration of the event (Fig. 4D, Movie S1). With the removal of sorbitol, the proportion of cells with WDR62 granules decreased rapidly (Fig. 4E, F), with a reduction in granule number and proportion of cells with granules evident as early as 10 min recovery and was significant by 30 or 60 mins recovery from sorbitol treatment (Fig. 4F-G). There was also a trend towards increased granule diameter with increasing recovery periods (Fig. 4H), which may reflect disassembly of smaller granules orthe sequential fusion of granules may occur during disassembly. When observed in real-time, we observed very rapid disassembly of WDR62 granules in cells within seconds of sorbitol removal (Fig. 4A). Thus, WDR62 granules appeared to be highly dynamic and underwent very rapid assembly/disassembly under permissive conditions.

**Figure 4.**
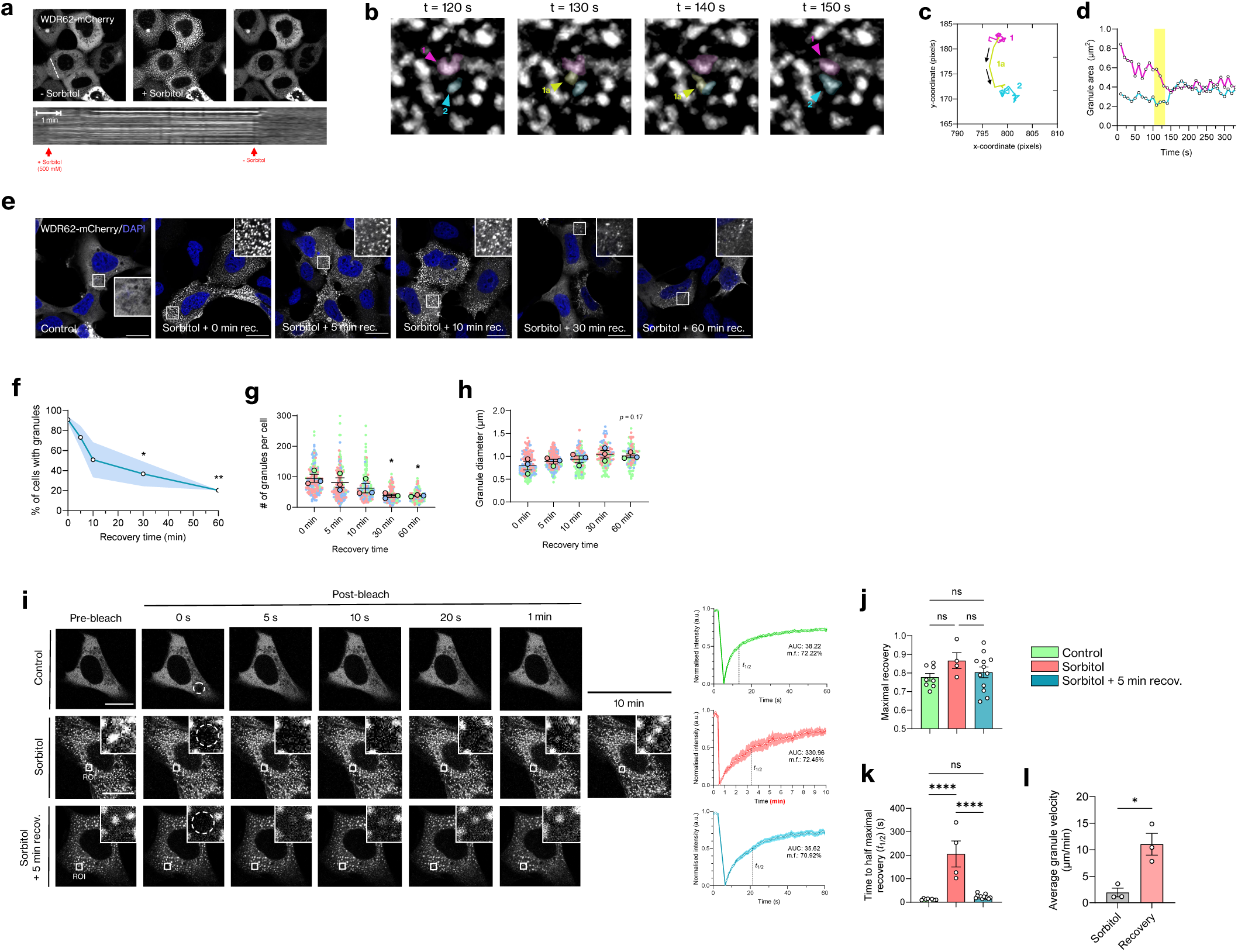
WDR62 granules are dynamic and liquid-like protein condensates. **(A)** Live-cell imaging experiments reveal that WDR62 granules assemble rapidly assemble (∼ 1 min) and disassemble (∼10 s) following the addition and removal of sorbitol stress. Bottom represents kymograph corresponding to white dashed line on first image. **(B-D)** WDR62 granules undergo fission and fusion events. **(E-H)** WDR62 granules disassemble following the removal of stress stimulus. One-way ANOVA with Tukey’s multiple comparisons (*p<0.05). **(I)** WDR62 granules exchange their contents with the bulk cytosol. *n* ≥ 4 granules. Shaded areas about the line represent SEM. Mobile fraction (m.f.) and area under the curve (AUC) displayed on graph. **(J)** Maximal fluorescence recovery as a fraction of pre-bleach intensity. One-way ANOVA with Tukey’s multiple comparisons (n.s. is p>0.05). **(K)** Time taken for bleached ROIs to recovery to half of the maximal level (*t*_1/2_). One-way ANOVA with Tukey’s multiple comparisons (****p<0.0001, n.s. is p>0.05). **(L)** WDR62 granule x-y velocity (µm/min). Recovery group is 5 min recovery following sorbitol washout. Two-tailed unpaired Student’s T-test (*p<0.05). All scale bars represent 20 µm, unless otherwise stated. Cell-level values for each quantification were separately pooled for each biological replicate, and the mean was calculated for each pool. Those three means were used to calculate the average (horizontal bar), the SEM (error bars), and P-values.

Liquid-like protein condensates also exchange their contents with the bulk cytosol [52]. Hence, we interrogated this behaviour in WDR62 granules with fluorescence recovery after photobleaching (FRAP) experiments. We quantified maximal fluorescence recovery (*F*_max_) relative to pre-bleach intensity and time to half maximal recovery (*t*_1/2_) for vehicle control, sorbitol-treated, and cells treated with sorbitol followed by 5 min recovery in iso-osmotic media. Maximal fluorescence recovery (*F*_max_) was similar across the three treatment groups with each recovering ∼80% of pre-bleached fluorescence (Fig. 4I, J). However, while diffuse WDR62 in control cells rapidly recovered from photobleaching (within 1 min), the rate of fluorescence recovery of WDR62 in granules of sorbitol-treated cells was significantly reduced (*t*_1/2_: 11.41 sec ± 1.298 vs. 205.8 sec ± 55.72, ****p≤0.0001) (Fig. 4I, K). We postulated that this reduction in WDR62 dynamics could be attributed to osmotic volume compression and reduced macromolecular diffusion caused by sorbitol treatment. Indeed, this was supported by a significant reduction in the dynamicity of WDR62 granules, as quantified by average granule velocity (µm/min) in the XY plane, in sorbitol treated cells compared to cells allowed to recover for 5 min in iso-osmotic media following sorbitol washout (recovery) (1.95 ± 0.82 vs. 11.03 ± 2.05, *p<0.05) (Fig. 4L). Consistent with this, the fluorescence recovery of WDR62 granules in the recovery phase was much more rapid and similar to diffuse WDR62 in control cells (*t*_1/2_: 23.2 ± 2.5 vs. 11.41 ± 1.298, n.s.) (Fig. 4I-K). Thus, WDR62 granules are highly dynamic and rapidly exchange with bulk cytosol when not constrained by hyperosmolarity.

The phase separation of proteins is commonly driven by weak, multivalent protein interactions including non-specific binding involving intrinsically disordered regions (IDRs) [53]. Indeed, according to disorder structure predictions, the C-terminal half of WDR62 is almost entirely disordered (Fig. S6A). To determine which WDR62 domains were involved in granule assembly, we expressed EGFP-tagged truncation mutants lacking the C-terminal disordered region (WDR62(N)) or the N-terminal WD repeat region (WDR62(C)) in AD293 cells (Fig. S6B). In response to sorbitol, WDR62(C) nucleated granules in almost all (∼95%) cells, similar to full-length WDR62 (WDR62(FL)) (∼92%), while WDR62(N) showed markedly reduced granule assembly (∼33%) (Fig. S6C-E, I). This indicated that sorbitol-induced WDR62 condensation was driven largely by interactions mediated by its intrinsically disordered C-terminal region. In addition to IDRs, the formation of protein condensates is mediated by multivalency achieved by interactions between protein dimerization or oligomerization domains [26]. Therefore, we next investigated the role of the C-terminal helix-loop-helix motif (dimerisation domain)—which is involved in the homo- and heterodimerisation of WDR62 [33]—on the formation of WDR62 granules. We utilised a truncated mutant of WDR62 which lacks the dimerisation domain (WDR62(1-1329)) (Fig. S6B) and noted a marked reduction in granule assembly in comparison to full-length WDR62 (Fig. S6F, I). To determine if the dimerisation domain could independently induce WDR62 condensation, we used two myc-tagged WDR62 truncation mutants comprised of the C-terminal domain without the dimerisation domain (WDR62(842-1329)) or one containing only the dimerisation domain (WDR62(1290-1523)) (Fig. S6B), neither of which assembled granules following sorbitol treatment (Fig. S6F-H). Together, this suggests that the disordered C-terminal region and dimerisation domains were both required for sorbitol-induced WDR62 condensation.

### WDR62 co-localizes with enzymes and co-chaperones involved in purine metabolism

We next explored WDR62 co-localisation with the HSP co-chaperones identified in our proximity labelling studies by co-expressing EGFP-tagged BAG2, STIP1 or DNAJC7 with WDR62-mCherry in AD293 cells. We found a large degree of overlap between BAG2 and WDR62, particularly in cytoplasmic granules assembled following sorbitol treatment (Fig. 5A). Similarly, STIP1 and DNAJC7 were also co-located in WDR62 granules formed in response to osmotic stress (Fig. 5B, C). The vast majority of WDR62 granules were comprised of these co-chaperones and their overlap with WDR62 fluorescence was substantial with an average Pearson’s correlation coefficient (PCC) of 0.9, 0.79 and 0.95 for BAG2, STIP1 and DNAJC7, respectively (Fig. 5H).

**Figure 5.**
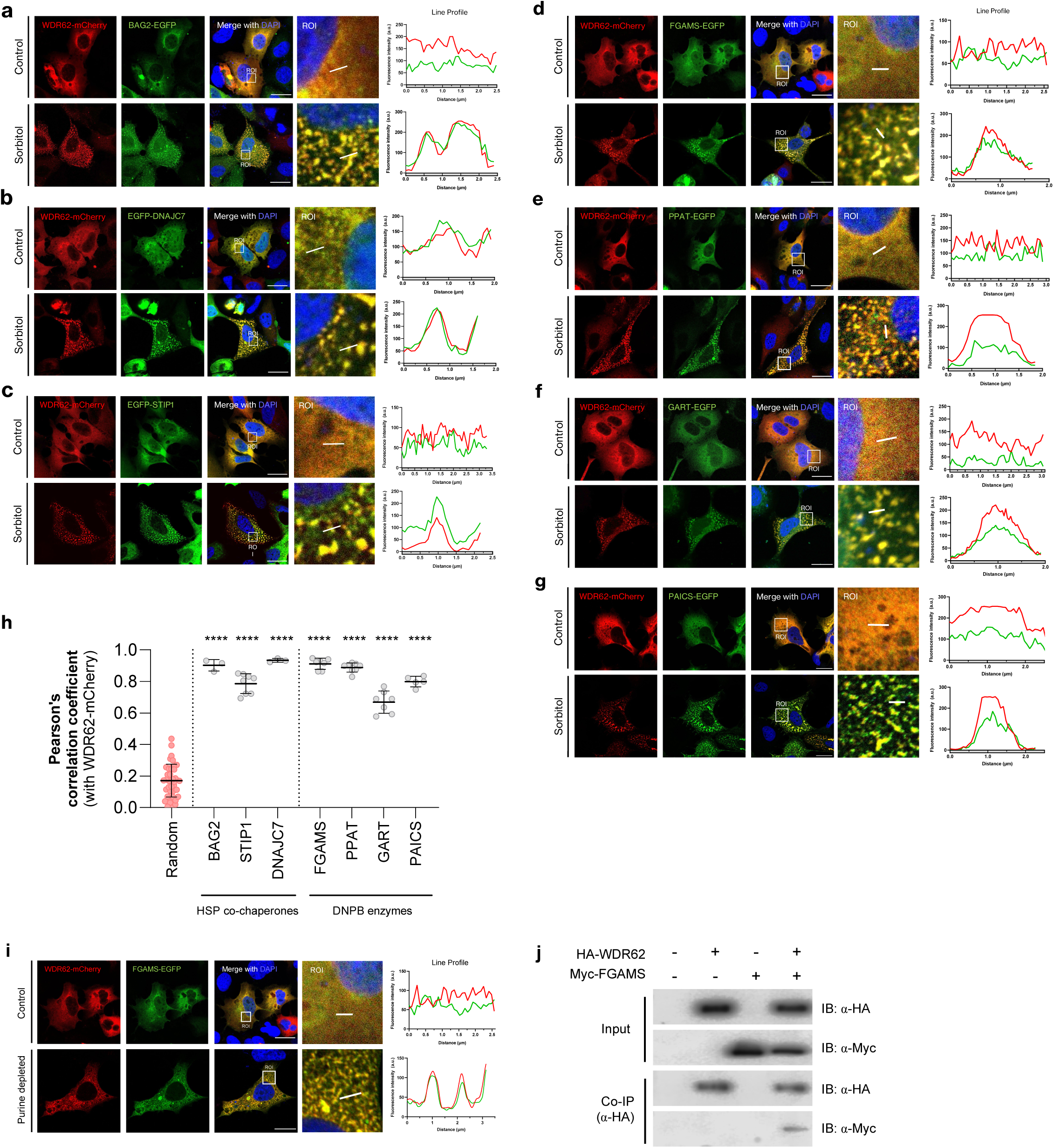
Members of the HSP70/HSP90 chaperone family and *de novo* purine biosynthesis pathway localise to WDR62 granules. Representative confocal micrographs of AD293 cells treated with 0.5 M sorbitol for 1 h. Cells transiently transfected with WDR62-mCherry and either the HSP co-chaperones **(A)** BAG2-EGFP, **(B)** EGFP-DNAJC7, **(C)** EGFP-STIP1, or *de novo* purine biosynthesis (DNPB) enzymes **(D)** FGAMS-EGFP**, (E)** PPAT-EGFP, **(F)** GART-EGFP or **(G)** PAICS-EGFP. Co-localisation of each signal is indicated by fluorescence intensity plots to the right of each set of images (y-axis represents fluorescence intensity (a.u.), x-axis (µm) represents the length of the white line drawn on the ROI). **(H)** Scatterplot representing the co-localisation between WDR62-mCherry and EGFP signal for each respective protein (mean ± SD). Each dot represents the Person’s correlation coefficient for a single ROI. One-way ANOVA with Tukey’s multiple comparisons (****p<0.0001). Data represent n = 3 biological replicates. Scale bars represent 20 µm. **(I)** Confocal micrographs of AD293 cells co-transfected with WDR62-mCherry and FGAMS-EGFP, in either control (top) or purine depleted (bottom) conditions. **(J)** Co-immunoprecipitation and immunoblot of myc-FGAMS and HA-WDR62.

HSP70/90 chaperones and co-chaperones—including BAG2, STIP1 and DNAJC7—co-cluster and are involved in the assembly of purinosomes, biomolecular condensates of metabolic enzymes involved in purine metabolism [25]. Therefore, WDR62 interaction and colocalization with these molecular co-chaperones suggests that WDR62 granules may represent purinosomes. We demonstrated that metabolic enzymes involved in *de novo* purine biosynthesis (DNPB) and assembled into purinosomes, such as FGAMS (also referred to as PFAS), PPAT, GART and PAICS, were all colocalized with WDR62 following sorbitol treatment (Fig. 5E-G). The fluorescence of these purine enzymes tagged with EGFP correlated near perfectly with mCherry-tagged WDR62 (Fig. 5H). Moreover, we showed that WDR62 also localized to *bona fide* purinosomes induced by purine-depleted media, independent of sorbitol treatment (Fig. 5I). Thus, WDR62 was assembled into multicomponent structures that contain DNPB enzymes and their associated chaperone machinery. In support of this, we show that HA-tagged WDR62 and myc-tagged FGAMS can be co-immunoprecipitated, suggesting they form a complex with BAG2 and related co-chaperones (Fig. 5J). From these studies, we concluded that WDR62 granules represent or overlap substantially with the phase-separated metabolons known as purinosomes.

### WDR62 loss sensitizes cells to purine depletion and dysregulates purine metabolic pathways

Our finding that chaperones and purine pathway enzymes were co-located in WDR62 granules raised the question of the role of WDR62 in the regulation of purine metabolism. Compared to wild-type (WT) AD293 cells, the CRISPR/Cas9 deletion of WDR62 (WDR62 KO) did not alter the formation, number, or morphology of purinosomes induced by sorbitol treatment or by purine-depletion (Fig. S7A-C). Further, WDR62 is a microtubule-associated protein [5] and microtubule-directed transport is involved in purinosome assembly [54]. However, WDR62 loss did not overtly change the close association of purinosomes with microtubules or mitochondria (Fig. S8). As WDR62 did not appear to be required for the formation of purinosomes, we next moved to determine if WDR62 was required for purine synthesis and purine-dependent cell growth/survival.

First, we examined the viability of WT cells compared to WDR62 KO cells when cultured in purine-depleted media. An immunoblot analysis confirmed the complete loss of WDR62 expression in KO cells (Fig. 6A). With purine-depletion, we observed a significantly higher percentage of rounded WDR62 KO cells compared to WT at 4 (31.83 ± 3.31% vs 12.82 ± 2.04, **p≤0.01) and 7 days (48.99 ± 1.64% vs 23.54 ± 2.67, **p≤0.01) in culture (Fig. 6B). In purine-rich (basal) conditions, there was no appreciable difference between WT and KO cells (Fig. 6B). Hence, the proliferation of WDR62 KO cells was significantly reduced after 4 and 7 days culture in purine depleted media compared to WT cells (Fig. 6C). Similarly, an assessment of LDH release as a measure of cell toxicity revealed a significant increase in WDR62 KO cells in purine-depleted conditions compared to WT cells (41.08 ± 7.45% vs 17.89 ± 1.87%, Fig. 6D). This accounted for a ∼2.4-fold increase in cytotoxicity in KO cells in purine depleted conditions, (**p≤0.01) whereas there was no significant difference between WT and KO cells under purine-rich conditions.

**Figure 6.**
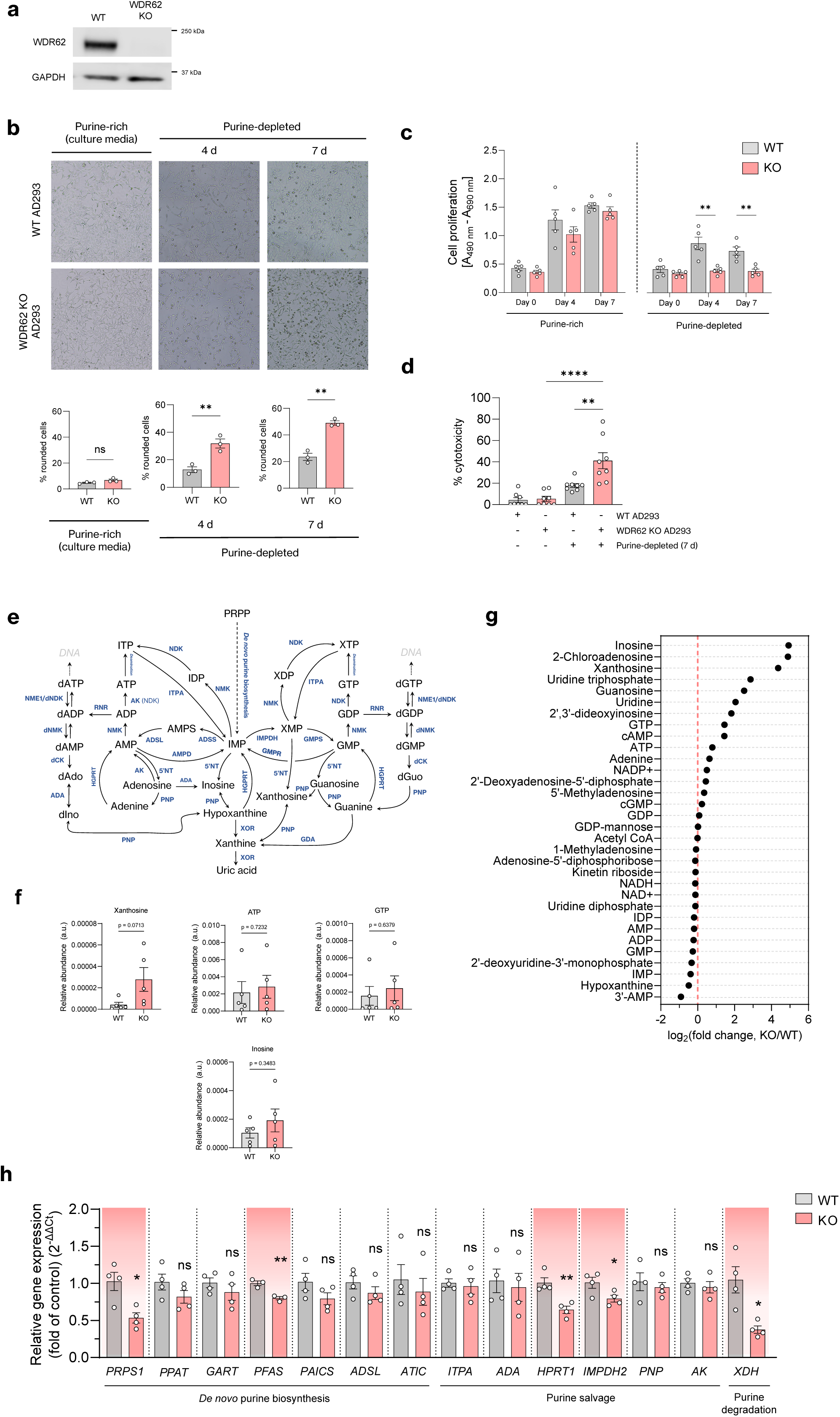
WDR62 regulates cellular purine metabolism. **(A)** Western blot confirming the loss of WDR62 expression in WDR62 KO cells. **(B)** Representative images of WT and WDR62 KO cells cultured in purine-rich or purine-depleted media. Bar graphs depict proportion (%) of rounded cells in each category. Two-tailed unpaired T-test. **(C)** XTT assay quantification of cell proliferation (A_490nm_-A_690nm_) of WT and KO cells cultured for 0, 4 and 7 days in purine-rich or purine-depleted media. *n* = 5 biological replicates. (**D)** LDH assay of percentage cytotoxicity (% cytotoxicity) of WT and KO cells cultured for 7 days in purine-rich or purine-depleted media. *n* = 8 biological replicates. Data in (B, C) analysed by one-way ANOVA with Tukey’s multiple comparisons (**p≤0.01, ****p≤0.0001, n.s. is p>0.05). **(E)** Schematic representation of cellular purine metabolism. **(F)** LC-MS/MS analysis of metabolite extracts harvested from WT and WDR62 KO cells grown in purine-depleted media for 1 week. Relative abundance of representative metabolites related to purine metabolism (n = 5 biological replicates). a.u. is arbitrary units. Data are shown as mean ± SEM. Two-tailed unpaired T-tests used for statistical analysis. Exact P-values are indicated. **(G)** Graph represents the differential metabolites between WT and KO cells (log_2_(fold change, KO/WT)) (n=8). **(H)** qPCR of WDR62, de *novo* purine biosynthesis (*PRPS1, PPAT, GART, PFAS, PAICS, ADSL, ATIC)* purine salvage (*ITPA, ADA, HPRT1, IMPDH2, AK)* and purine degradation (*XDH)* gene expression in WT and WDR62 KO AD293 cells. Asterisks denote statistical significance (two-tailed unpaired T-test) in comparison of WT and KO groups for each target gene. *n* = 4 biological replicates.

Investigating how WDR62 depletion alters purine synthesis pathways (Fig. 6E), we again upregulated *de novo* purine synthesis by culturing cells in purine-depleted media and analysed metabolite extracts from WDR62 KO cells compared to WT with liquid-chromatography tandem mass-spectrometry (LC-MS/MS). While the average relative abundance of AMP, GMP and IMP levels were comparable between WT and KO cells, our analysis revealed an accumulation of purine nucleotide intermediates such as inosine, xanthosine, guanosine and analogues of adenosine such as 2-chloroadenosine (Fig. 6F, G). We next investigated the mRNA expression of a panel of enzymes involved in purine metabolism by qPCR. In WDR62 KO cells, we found a significant downregulation of enzymes involved in *de novo* purine synthesis such as *PRPS1* (∼48% reduction) and *PFAS* (∼20% reduction), purine salvage genes *HPRT1* (∼39% reduction) and *IMPDH2* (∼22% reduction), and the purine degradation gene *XDH* (∼64% reduction) (Fig. 6H). Taken together, our findings indicate that WDR62 loss led to increased cell sensitivity to purine depletion and dysregulation of purine metabolic pathways.

### WDR62 regulates the expression and stability of HPRT

In our metabolomic analysis, the accumulation of purine nucleosides, such as inosine, xanthosine and guanosine suggests WDR62 depletion led to altered recycling of purine bases, rather than a defect in purine synthesis. Therefore, we focused our study on the purine recycling enzyme HPRT and confirmed that protein levels were substantially reduced (∼49%) in WDR62 KO cells (Fig. 7A) and this was rescued by exogenous expression of WDR62-GFP in KO cells (Fig. 7B). In contrast, despite reduced mRNA (Fig. 6H), the protein levels of the PFAS enzyme involved in *de novo* purine synthesis were not appreciably different between WT and WDR62 cells (Fig. 7A). While the reduction in *HPRT* mRNA levels in WDR62 KO cells is consistent with transcriptional downregulation (Fig. 7A), the identified proximity partners of WDR62 suggests roles regulating protein folding or stability (Fig. 1). To determine if post-translational mechanisms also contributed to WDR62-dependent loss of HPRT, we performed a cycloheximide (CHX) chase experiment to evaluate the half-life of HPRT (Fig. 7C). Following CHX inhibition of protein synthesis, we found that the degradation of HPRT was much more rapid in WDR62 KO cells compared to WT (Fig. 7C). However, the degradation of the purine synthesis enzyme PFAS was not appreciably increased in WDR62 KO cells (Fig. 7C). Thus, WDR62 is required to maintain the stability of HPRT.

**Figure 7.**
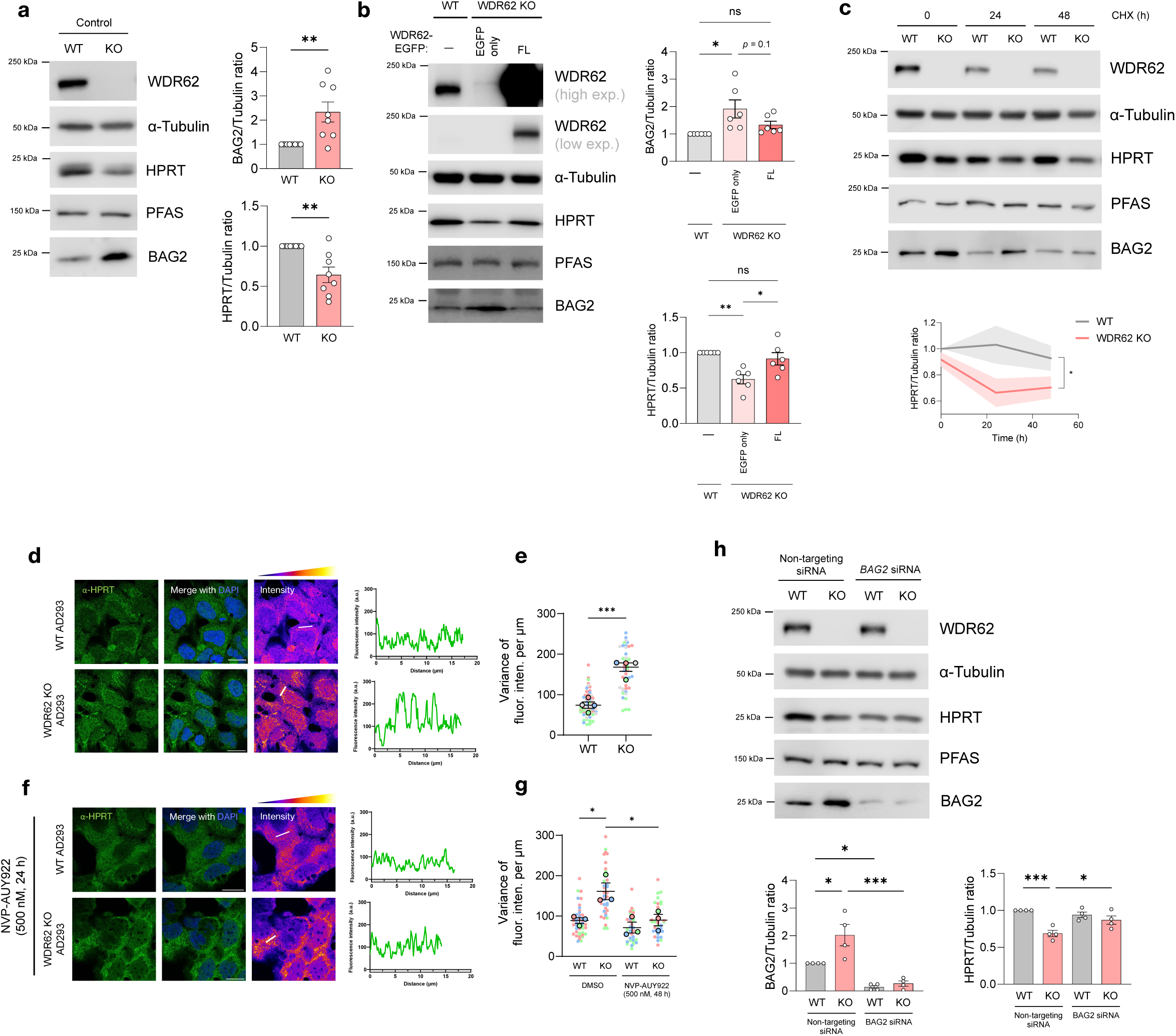
The loss of WDR62 drives a BAG2-dependent loss of purine metabolic enzymes. **(A)** Immunoblot of purine metabolic enzymes PFAS, HPRT and XDH and co-chaperones BAG2 and DNAJC7 in WT and WDR62 KO AD293 cells. **(B)** Immunoblot of WDR62 KO AD293 cells rescued with EGFP only or WDR62(FL)-EGFP. **(C)** Cycloheximide chase assay examining the half-life of HPRT, PFAS and BAG2 in WT and WDR62 KO cells. WT and WDR62 KO cells treated with 100 µM CHX for 0, 24, or 48 h. **(D)** WT AD293 cells treated with non-targeting or *WDR62* siRNA and immunoblot for BAG2, HPRT and PFAS. **(E)** Western blot analyses of PFAS, HPRT and BAG2 in WT AD293 cells transfected with EGFP only or WDR62-EGFP. **(F, G)** Immunofluorescence and confocal micrographs of endogenous HPRT in vehicle (DMSO) or NVP-AUY922-treated WT and WDR62 KO cells. Far right image represents pseudo-coloured “fire” LUT to better visualise pixel intensity. Fluorescence intensity plots to the right of each set of images (y-axis represents fluorescence intensity (a.u.), x-axis (µm) represents the length of the white line drawn on pseudo-coloured intensity image. **(E, G)** SuperPlots of averaged normalised (per µm) variance of fluorescence intensity along random lines plotted in the cytoplasm of random cells. Higher variances values indicate greater heterogeneity in fluorescence intensity, characteristic of a less diffuse and more punctate distribution. **(H)** Western blot analyses of WDR62, BAG2, HPRT and PFAS expression in WT and WDR62 KO cells treated with non-targeting or *BAG2* siRNA.

In the absence of WDR62, we also observed altered subcellular localisation of endogenous HPRT through immunofluorescence (Fig. 7D). Notably, HPRT expression in the cytoplasm of WDR62 KO cells was less diffuse and exhibited a more condensed localisation (Fig. 7D). In the absence of other cell treatments, HPRT did not form distinct purinosome-like puncta. In a line scan analysis, HPRT fluorescence exhibited greater signal variance (normalised to line length) in WDR62 KO cells compared to WT cells, which suggests increased propensity for HPRT to aggregate when WDR62 is lost (Fig. 7E). Since HSP90 is known to regulate the stability and folded state of many purine metabolic enzymes [24, 26], we treated cells with a small molecule inhibitor of HSP90, NVP-AUY922. Interestingly, this reversed the condensed morphology of HPRT in WDR62 KO cells (Fig. 7F, G). This indicates that the stability and cytoplasmic distribution of HPRT was regulated by HSP90, and this was associated with WDR62.

### Cellular HPRT levels are regulated by WDR62-BAG2

BAG2 is a co-factor for molecular chaperones such as HSP70 but is also associated with HSP90 [55]. In some pathological contexts, BAG2 has been reported to promote the misfolding and aggregation of cytoplasmic proteins in a HSP90 dependent manner [56]. Given the interaction between WDR62 and BAG2 (Fig. 2), we next investigated BAG2 levels and observed a ∼2.6-fold increase in BAG2 expression in parallel with reduced HPRT levels following the loss of WDR62 (Fig. 7A). This effect was replicated by transient depletion of WDR62 with siRNA (Fig. S9A) and rescued with WDR62-EGFP in WDR62 KO cells (Fig. 7B). Conversely, the overexpression of WDR62 in WT cells led to a significant decrease in BAG2 levels (Fig. S9B), indicating that WDR62 negatively regulates cellular levels of BAG2. Intriguingly, HPRT levels were reduced with WDR62 overexpression in WT cells (Fig. S9B). So, while enhanced BAG2 levels were associated with downregulated HPRT in absence of WDR62, a decrease in BAG2 expression from ectopic overexpression of WDR62 does not necessarily lead to increased HPRT expression. Unlike BAG2, the expression of other chaperone interactors of WDR62 that were identified in our BioID study and co-located in WDR62 granules (Fig. 1) (Fig. 5A-C), such as HSP70, HSP90, and co-chaperones STIP1 and DNAJC7, remained unchanged (Fig. S9C).

Subsequently, we targeted BAG2 with siRNA to determine if elevated BAG2 in WDR62 KO cells contributed to the diminished half-life of HPRT. Compared to WT cells, HPRT levels were significantly reduced in WDR62 KO cells transfected with non-targeting siRNA, as expected (Fig. 7H). Importantly, siRNA depletion of BAG2 in WDR62 KO cells restored HPRT to WT levels (Fig. 7H). Taken together, our studies revealed that the WDR62-BAG2 axis was required to maintain HPRT stability and the proper folding and cytosolic distribution of HPRT in a HSP90-dependent manner. These new mechanisms contribute to dysregulation of purine metabolism which may underlie the development of congenital microcephaly associated with *MCPH2* mutations.

## Discussion

Studies to date have primarily focused on mitosis and centriole biogenesis as roles for WDR62 in neurodevelopment. Our study presents evidence of a novel role for cytoplasmic WDR62 and the BAG2 co-chaperone in a regulatory network that governs the stability and turnover of purine metabolic enzymes, thereby controlling purine homeostasis. We also show that WDR62 forms dynamic, phase-separated granules that co-localise with chaperones and purine metabolic enzymes, resembling purinosomes. Although WDR62 was not required for purinosome assembly, we demonstrated that the WDR62-BAG2 axis was involved in regulating cellular purine homeostasis and specifically the stability of the purine salvage enzyme, HPRT.

Our studies indicate that BAG2 interacts with WD-repeat region and the C-terminal α-helical dimerization domain of WDR62. Furthermore, we show that microcephaly-associated mutations alter WDR62-BAG2 interactions, suggesting that defects in this novel regulatory pathway of purine metabolism may contribute to abnormal neurodevelopment. However, it’s unclear why microcephaly-linked missense mutations in WDR62 enhanced BAG2 interactions despite not being contained within the predicted WDR62-BAG2 interface (Fig. 2). It is possible that these specific residues might participate in uncharacterised protein-protein interactions that normally sterically impede on the nearby BAG2 binding site, thereby potentially enhancing WDR62-BAG2 interactions when mutated. Conversely, we found that frameshift mutations such as 3936dupC, which truncate the C-terminal dimerisation domain while preserving the full N-terminus, disrupt binding to BAG2 (Fig. 2E). This suggests that although the N-terminal half can engage with BAG2, the disordered C-terminus may impede the WDR62-BAG2 dimer in the absence of the dimerisation domain. Supporting this idea, we observed a trend towards increased WDR62(N) interaction with BAG2 compared to full-length WDR62 (Fig. 2D).

Our findings that microcephaly-associated WDR62 mutations can lead to increased or decreased binding of BAG2, depending on the specific mutation. This suggests that either excessive or deficient interactions between WDR62 and BAG2 may be detrimental. Consistent with this idea, WDR62 and BAG2 expression levels were feedback regulated, where the loss of WDR62 leads to an increase in BAG2 and vice versa (Fig. 7A) (Fig. S9B). We speculate that V65M and R438H missense mutations and increased WDR62-BAG2 interactions may sequester BAG2, disrupting its normal function. In contrast, frameshift mutations may reflect a knockout effect with decreased BAG2 binding and regulation. Hence, an abnormal increase or decrease in WDR62-BAG2 interactions may similarly result in disruptions in purine enzyme turnover and purine homeostasis.

WDR62 forms cytoplasmic granules which co-localise with chaperone machinery and purine metabolic enzymes. These granules resembled purinosomes and we described their assembly in response to hyperosmotic stress, which triggers protein phase separation through mechanisms such as cell volume exclusion and macromolecular crowding [57, 58]. However, it is unclear if WDR62 granules contain all necessary components of the *de novo* purine biosynthesis (DNPB) pathway and/or represent functional metabolons which enhance metabolic flux under conditions of purine depletion, hypoxia, or casein kinase 2 (CK2) inhibition [47, 59, 60]. In contrast, there is evidence of metabolically-inactive purine enzyme assemblies that sequester pathway components, hindering *bona fide* purinosome assembly and metabolic flux [26]. Nonetheless, our study confirms that WDR62 granules co-localise with at least four of six DNPB enzymes (Fig. 5D-G), and we observed presence of WDR62 within *bona fide* purinosomes formed under purine depletion conditions (Fig. 5I). However, while HSP70/90, their co-chaperones, and other WD repeat-containing proteins are involved in purinosome assembly [61], we show WDR62 is not essential for this process (Fig. S7). Consistent with this, our metabolite analysis did not indicate overt defects in de novo purine biosynthesis, but instead the accumulation of purine intermediates caused by altered nucleotide recycling (Fig. 6F-G). Therefore, while we revealed a role for WDR62 in maintaining purine homeostasis, the significance of WDR62 located in purinosome granules remains undetermined.

The mechanism behind BAG2-mediated HPRT degradation after WDR62 loss (Fig. 11) remains to be fully elucidated. Indeed, low stoichiometric concentrations of BAG2 stimulates ATP turnover and enhances chaperone cycling of HSP70-DNAJ pairs [62]. Conversely, high concentrations of BAG2 inhibit ATP turnover, stabilise unbound HSP70, and inhibit folding activity [62]. Thus, BAG2 appears to have a dual role in proteostasis, inhibiting protein folding of HSP70/90—and possibly enhancing degradation—when expressed at high levels, and with pro-folding roles at lower levels. Thus, increased BAG2 expression following WDR62 loss may be a compensatory mechanism to enhance the degradation of misfolded clients. Indeed, BAG2 promotes the degradation of large protein aggregates, such as phosphorylated Tau, via ubiquitin-independent proteasomal pathways [38, 63], and activates autophagy in protein aggregation contexts [64, 65]. Consistent with this, we revealed that endogenous HPRT expression is also less uniformly distributed in WDR62 KO cells, suggesting a loss of HPRT solubility (Fig. 7C). Notably, inhibition of HSP90 restored the normal distribution of HPRT, indicating a role for HSP90 in proper HPRT folding and stability (Fig. 7E). In line with this, elevated BAG2 expression has been shown to promote the aggregation of mutant p53, rescued by BAG2 knockdown or inhibition of HSP90 [56]. Further, HSP90 is also involved in regulating the solubility of FGAMS [26].

However, the specific mechanism underlying the enhanced degradation of HPRT in WDR62 KO cells is unknown. Indeed, many purine metabolic enzymes are clients for proteasomal degradation [24]. Interestingly, another WD repeat protein linked to microcephaly, WDR47, regulates proteostasis by controlling autophagic flux [66]. However, the role of WDR62-BAG2 in maintaining the stability of purine enzymes may also extend beyond post-translational mechanisms and to transcriptional regulation. This is supported by our observation of decreased *HPRT* mRNA levels in WDR62 KO cells (Fig. 6H). Indeed, BAG family proteins can exhibit transcription factor activity and can modulate the transcriptional activity of various clients, including those related to nuclear hormone receptor activity [67, 68]. Further, purine intermediates like AICAR and SAICAR can also influence dimeric interactions between transcription factors, thereby regulating purine homeostasis at the transcriptional level [69]. Hence, the downregulation of purine pathway genes in WDR62 KO cells may also originate from feedback mechanisms due to accumulation of purine intermediates, caused by an initial loss of BAG2-regulated pathway enzyme stability.

It remains to be determined whether the purine metabolic defects observed in our in vitro experiments would be replicated by WDR62 mutations in patients, and which tissues or cell types would exhibit the most pronounced defects. Purines play a role in supporting the rapid growth, division, and differentiation of neural progenitors during neurogenesis by providing nucleotides, energy, and purinergic signalling, all of which are all crucial for proper brain development [70–73]. In line with this, the loss of purine metabolic enzymes, including HPRT, disrupts neurogenesis, resulting in microcephaly, cell cycle defects, ciliopathies, and abnormalities in proliferation and neural progenitor fate decisions, mirroring the loss of WDR62 [29]. Moreover, other WD repeat-containing and microcephaly-associated proteins, including NWD1, interact with purine enzymes such as PAICS to facilitate purinosome assembly, ensuring proper neural differentiation and migration [61]. While specific roles for chaperone machinery in neurodevelopment are established, they have not been specifically identified for BAG2 [74]. Overall, our study has uncovered a novel role for WDR62-BAG2 in maintaining HPRT stability and purine metabolism, which may contribute to neurodevelopmental deficits in *MCPH2* patients. Given purine metabolic dysfunction can cause neural stem cell defects [75], it would be of interest to determine whether WDR62 regulation of purine enzymes represents the primary defect contributing to abnormal brain growth in this context.

## Materials and Methods

### Antibodies, reagents, and plasmids

Primary antibodies used in this study are outlined in detail in Table S1. Secondary antibodies used in this study were: anti-rabbit IgG (H+L) Alexa Fluor 488 (Invitrogen, A11008, 1:800 (IF)); anti-mouse IgG (H+L) Alexa Fluor 555 (Invitrogen, A21422, 1:800 (IF)); goat anti-rabbit IgG (H+L) HRP (Invitrogen, 31460, 1:10000 (WB)); goat anti-mouse IgG (H+L) HRP (Invitrogen, 62-6520, 1:10000 (WB)); donkey anti-goat IgG (H+L) HRP (Invitrogen, A15999, 1:5000 (WB)).

Origins of all plasmids used in this study outlined in Table S2. Traditional restriction enzyme cloning (double-restriction digestion and ligation) techniques were used for all cloning procedures. The empty pmCherry-N1 and pmCherry-C1 vectors were purchased from Clontech. Empty pXJ40-myc, -HA and -FLAG vectors are from [76]. Recombinant plasmids formed with the above vectors was completed using standard restriction-enzyme cloning techniques with PCR-amplified inserts generated with specific primers. Plasmids ligated using the Rapid DNA Ligation Kit (Thermo Scientific, K1422) as per manufacturer’s instructions. All constructs validated by full sequence analysis. Primers outlined in Table S3.

### AlphaFold2 protein structure prediction

Amino acid sequences for human WDR62 and BAG2 were retrieved from UniProt. The ColabFold v1.5.5 implementation of AlphaFold2 was used [77] on a cloud 8x Tesla V100 16GB GPU. PAE Viewer (http://www.subtiwiki.uni-goettingen.de/v4/paeViewerDemo) was used to view predicted structure and predicted alignment error (PAE). Proteins, Interphases, Structures and Assemblies (PDBePISA) (https://www.ebi.ac.uk/pdbe/pisa/) was used to calculate interphase area (Å) for predicted complexes.

### Cell culture, transfections, and chemicals

WT AD293 cells (WT) (Agilent Technologies, Inc.) and AD293 cells deleted of WDR62 by CRISPR/Cas9-sgRNA (WDR62 KO) were maintained by standard protocols [78]. In brief, cells were maintained in culture media containing Dulbecco’s Modified Eagle Medium (DMEM) (Gibco, 21969035) supplemented with 10% (v/v) foetal bovine serum (FBS) (Gibco, 10099141) and 100 µg/ml penicillin-streptomycin (Pen-Strep) (Gibco, 15140122). Cells cultured at 37°C and 5% CO_2_ within a humidified incubator. DNA transfections were performed with Lipofectamine™ 2000 (Invitrogen, 11668019) according to the manufacturer’s instructions. Where indicated, cells were treated with sodium arsenite (arsenite; 0.5 mM; Sigma-Aldrich, S7400), D-sorbitol (sorbitol; 0.5 M; Sigma-Aldrich, S3889); sucrose (0.5 M; Chem-Supply, SA030), dextrose (0.5 M; Sigma-Aldrich, D9439); cycloheximide (CHX; 100 µg/ml; Sigma-Aldrich, 01810), NVP-AUY922 (NVP; 500 nM; Cayman Chemical, 10012698), sodium chloride (NaCl; 0.5 M; Sigma-Aldrich, J21618); betaine (20 mM; Sigma-Aldrich, 61962); MG-132 (MG132; 10 µM; Abcam, ab141003) and; chloroquine diphosphate salt (CQ; 50 µM; Sigma-Aldrich, C6628).

### RNAi

RNA transfections were performed using Lipofectamine™ RNAiMAX (Invitrogen, 13778150) according to manufacturer’s instructions. ON-TARGETplus non-targeting control pool (D-001810-10) was used as the siRNA control. ON-TARGETplus human *BAG2* pool (L-011961-00), and human *WDR62* pool (L-031771-01) were obtained from Dharmacon™ (Horizon Discovery).

### Immunofluorescence

Cells grown on square 22 mm^2^ uncoated glass coverslips in 6-well plates. All steps performed at room temperature unless otherwise stated. Cells rinsed three times quickly with ice-cold PBS, then fixed with 4% (w/v) paraformaldehyde (PFA) in PBS for 20 min. Cells rinsed three times in PBS for 5 min each, then permeabilised with 0.2% (v/v) Triton X-100 in PBS (PBS-T) for 10 min. Cells rinsed again three times in PBS for 5 min each, then blocked with 1% (w/v) bovine serum albumin (BSA) (Sigma-Aldrich, A9418) in PBS-T for 30 min. Cells subsequently stained with appropriate primary antibodies (Table S1), and then with appropriate fluorophore-conjugated secondary antibodies for 1 h in darkness. Unless otherwise stated, antibodies diluted in 1% BSA in PBS-T. Cells rinsed three times in PBS for 5 min each, counterstained with DAPI for 5 min, and mounted in ProLong™ Diamond Antifade Mountant (Invitrogen, P36970) on glass slides. Cells overexpressing fluorescently tagged protein constructs were simply fixed, washed and counterstained with DAPI (Invitrogen, D1306).

### Protein extractions, immunoblots and co-immunoprecipitation

After treatments, cells were lysed with RIPA buffer [150 mM NaCl, 100 mM Na_3_VO_4_, 50 mM Tris-HCl pH 7.3, 0.1 mM EDTA, 1% v/v Triton X-100, 1% w/v sodium deoxycholate and 0.2% w/v NaF] supplemented with protease inhibitors (cOmplete™, Mini, EDTA-free Protease Inhibitor Cocktail, Roche, 11836170001). Protein lysates were cleared by centrifugation (16000 *g*, 15 min) and protein concentrations determined by protein assay with Bradford dye reagent (Bio-Rad, 5000006). Proteins separated by SDS-PAGE without boiling due to the heat sensitivity of WDR62 [7], then transferred onto polyvinylidene fluoride (PVDF) membranes and immunoblotted.

For co-immunoprecipitation experiments, cleared cell lysates were incubated with rabbit anti-HA primary antibodies, followed by affinity isolation with Protein A agarose beads (Roche, PROTAA-RO) overnight at 4°C on an end-over-end rotator. Beads pelleted and washed thoroughly in RIPA buffer with fresh protease inhibitors. Immunoprecipitated proteins eluted from beads with Laemmli buffer and separated by SDS-PAGE for immunoblotting.

For GFP-Trap immunoprecipitation experiments, cell lysates were incubated with GFP-Trap® agarose (ChromoTek, gta) for 1 hour at 4°C on an end-over-end rotator. Beads subsequently pelleted and washed thoroughly in RIPA buffer. Immunoprecipitated proteins eluted with Laemmli buffer and separated by SDS-PAGE for downstream immunoblot.

### Micrograph acquisition and processing

Immunofluorescence and fluorescence images were acquired with a Leica SP8 DMi8 inverted point scanning confocal microscope (Leica Microsystems, UK) equipped with a Leica HC PL APO 63x/1.30 GLYC CORR CS2 objective, or a Zeiss LSM900 inverted laser scanning confocal microscope with Airyscan 2 equipped with a 63x C-Plan Apo NA 1.4 oil-immersion objective. Image exposure times, gain and other settings were standardised between samples. Resolution was set at 1024 x 1024 DPI or 2048 x 2048 DPI with bi-directional scanning. Pinhole size set at 1. Image panels were assembled with ImageJ and Microsoft PowerPoint. Images collected using either Leica LAS X or Zeiss ZEN software and processed with ImageJ, where brightness and contrast were minimally adjusted across all panels indiscriminately. These adjustments merely enhanced viewing quality and did not obscure original information present in raw files.

### Live-cell imaging

Cells were seeded onto 8-well glass bottom chamber slides (Ibidi). Media was changed to DMEM, no phenol red, supplemented with 10% FBS and 100 µg/ml Pen-Strep 24 h prior to imaging. Imaging was routinely conducted on a Zeiss LSM900 inverted laser scanning confocal microscope, equipped with a 63x C-Plan Apo NA 1.4 oil-immersion objective and retrofitted with an incubation chamber to maintain physiological conditions (37°C and 5% CO_2_). Tracking of granules in time-lapse images conducted with the TrackMate plugin of ImageJ (Fiji).

### Fluorescence recovery after photobleaching (FRAP)

For fluorescence recovery after photobleaching (FRAP) experiments, the ZEISS ZEN FRAP tool was used to select a bleach area, which usually encompassed an individual granule, or a circular region of the cytoplasm ∼2 µm in diameter. Five pre-treatment images were taken, the area was bleached at 100% laser power for 100 iterations per ROI, and images were acquired every 5 seconds during recovery. Mean fluorescence intensity was analysed and graphed to an exponential recovery curve using ImageJ (Fiji).

### Cytotoxicity detection (LDH) assay

WT AD293 and WDR62 KO AD293 cells were seeded at 4 x 10^4^ cells/well in a 96-well tissue culture plate and cultured in purine-rich or purine-depleted conditions for 7 d. The next day, cells assayed for lactate dehydrogenase (LDH) activity with a Cytotoxicity Detection Kit^PLUS^ (LDH) (Roche, 11644793001) according to the manufacturer’s instructions. Absorbance (OD) measured on a FLUOstar Omega Microplate Reader (BMG Labtech) at 490 nm. Two controls were included: background control (cell-free, assay medium only) and high control (maximum LDH release, achieved by cell lysis in Triton X-100). To determine experimental absorbance values, average absorbance values of the sextuplicate samples and controls were calculated and absorbance values of background control were subtracted. Percent cytotoxicity was determined by the following equation: Cytotoxicity (%) = (exp. value/high control) x 100. Any negative values after subtracting average background absorbances were considered as zero.

### BioID assay and protein mass spectrometry

WT AD293 cells were seeded in 10 cm dishes, and transfected with WDR62-BirA*-HA, WDR62-HA or BirA*-HA the following day with Lipofectamine 2000, as per manufacturer’s instructions. The following day, cells were treated with 50 µM biotin for 24 h. Cells arrested in mitosis by low dose nocodazole treatment (200 ng/ml) represent *mitotic* populations. *Asynchronous* populations are untreated. Cells were washed in PBS and lysed in RIPA lysis buffer. Biotinylated proteins were captured with Pierce™ Streptavidin Agarose (Thermo Fisher Scientific, 20353) on a rotator for 4 h at 4°C. Agarose beads washed 3 times in RIPA buffer and were collected by centrifugation for 1 min between wash steps. For immunofluorescence, cells probed with Streptavidin-Cy3™ (Sigma-Aldrich, S6402). For western blot analysis, beads resuspended in Laemmli buffer with 1 mM biotin. Membranes probed with streptavidin-HRP or primary antibodies specific to proteins of interest. To identify proteins isolated by the BioID pulldown, biotinylated proteins were cleaved from streptavidin beads by tryptic digestion and were prepared for LC-MS/MS analysis.

### Cell proliferation (XTT) assay

WT AD293 and WDR62 KO AD293 cells were seeded at 2 x 10^4^ cells/well in a 96-well tissue culture plate and were incubated at 37°C and 5% CO_2_ in purine-rich or purine-depleted conditions. Cell proliferation was determined by XTT at Day 1, Day 4 and Day 7 after seeding. The XTT cell proliferation assay enables the quantification of cellular redox potential, providing a colorimetric readout of cell viability. Absorbance (OD) were obtained using a FLUOstar Omega Microplate Reader (BMG Labtech) at two wavelengths: 490 and 690 nm. The primary measurement wavelength was 490nm, corresponding to the peak absorbance of the formazan product. To account for non-specific background absorbance caused by culture media or other factors, absorbance at 690 nm was measured and subtracted from absorbance at 490 nm. Cell proliferation is then expressed as arbitrary units (a.u.) based on the subtracted value (A490nm-A690nm).

### Metabolite extraction and LC-MS/MS

Cells were seeded into T175 flasks and grown in purine-depleted media for 1 week. Cells were washed once in ice-cold PBS, harvested by scraping and thoroughly resuspended in 10 ml ice-cold PBS. A small sample (∼30 µl) was removed for cell counting on a hemocytometer. Cells were pelleted (200 *g,* 5 min, 4°C) and cell pellet was immediately quenched and resuspended in 800 µl of 100% methanol (pre-cooled at -80°C for ≥ 1 h). Quenched cell pellets were vortexed vigorously, thawed on ice for 10 min, then snap frozen in LN_2_ for 10 min. This freeze-thaw cycle was repeated two more times. Lysates centrifuged (7000 *g*, 15 min, 4°C) and supernatant collected. Pellet resuspended in 200 µl of 80% (v/v) methanol in water (also pre-cooled at -80°C) and extracted again by freeze-thaw as above. Combined supernatants from the two extractions were centrifuged (16000 *g*, 30 min, 4°C) to remove all insoluble material, dried with a Genevac™ miVac Quattro vacuum concentrator (7 mbar, ∼3 h, 35°C) and reconstituted in 30 µl of 80% (v/v) methanol in water with 0.1% (v/v) formic acid. Samples were then analysed by LC-MS/MS at the SCMB Mass Spectrometry Facility, The University of Queensland.

The LC platform consisted of an Exion UPLC system. Samples were separated on a Phenomenex Kinetex C18 column (100 mm x 2.1 mm, 1.7 µm particle size) using a water-methanol gradient. The UPLC column was maintained at 35°C and samples were stored in the autosampler at 4°C. Flow rate was 250 µl/min and injection volume was 10 µl. Solvent A was 0.1% formic acid in MilliQ water; solvent B was 100% methanol. The MS platform consisted of a SCIEX X500B QTOF mass spectrometer controlled by SCIEX Analyst software. The MS was operated in positive and negative ion mode and scanned from 6 to 1200 *m/z* in SWATH DIA (data independent acquisition) acquisition mode. Data were analysed using MS-DIAL software (ver 5.1.221218). MS-DIAL was used to match experimental mass spectra, corresponding to [M-H]- ions, against the MassBank mass spectral library. Compound identification was achieved by weighted similarity score after considering retention time, mass and MS/MS spectra. Ion identification was completed in MSFINDER. Relative abundance (a.u.) of metabolites was calculated by dividing the peak area (AUC) of each metabolite by the peak areas of all detected metabolites in the same sample (mTIC). mTIC represents the sum of all peak areas for ‘genuine’ or positively identified metabolites belonging to the same MS/MS scan, to avoid normalising against non-biological artifacts.

### RNA extraction, cDNA synthesis and qPCR analysis

Total RNA was extracted from WT and WDR62 KO AD293 cells using the PureLink™ RNA Mini Kit (Invitrogen, #12183020). 1 µg total RNA was used to perform reverse transcription using SuperScript™ III Reverse Transcriptase (Invitrogen, #18080044), as per manufacturer’s instructions. On-column DNase treatment was performed using the PureLink™ DNase Set (Invitrogen, #12185010). Real-time PCR was performed on a QuantStudio™ 7 Flex Real-Time PCR System (Applied Biosystems) after mixing cDNA, QuantiNova SYBR Green Master Mix (Qiagen, #208052) and gene-specific primers as listed in Table S4. Gene-specific primers were designed using the NCBI Primer-BLAST tool (https://www.ncbi.nlm.nih.gov/tools/primer-blast/), where all primer sets produced an amplicon 80-200 bp in size, had at least 2 mismatches within the last 5 bps at the 3’ end, and at least one primer spanned an exon-exon junction. Relative gene expression was calculated with the 2^-ΔΔCt^ method using GAPDH, β-actin and 18S rRNA as internal controls. Data was presented as fold-change relative to control (WT) cells.

### Analyses and Statistical Comparisons

All statistical analyses and the generation of graphs as conducted in GraphPad Prism 9 software (version 9.2.0). Densitometric analysis of Western Blot bands was performed in LI-COR Image Studio™ software. All bands normalised against a loading control (GAPDH or α-tubulin). The D’Agostino-Pearson test for normality was used to determine whether or not data assumed a Gaussian (normal) distribution. In groups which assumed Gaussian distributions, parametric tests were employed (e.g., two-tailed Student’s T-test, one-way ANOVA with Tukey’s multiple comparisons, and two-way ANOVA with Sidak’s multiple comparisons). In groups which did not assume a Gaussian distribution, non-parametric tests were employed (Mann-Whitney rank-sum test, Kruskal-Wallis test with Dunn’s multiple comparisons, and two-way ANOVA with Sidak’s multiple comparisons). The threshold of statistical significance was set at p≤0.05. All error bars represent mean ± standard error of mean (SEM). All figures were generated using GraphPad Prism 9, Microsoft PowerPoint, or Adobe Illustrator. Each experiment was conducted at least three times for at least three independent biological replicates (n=3).

For quantification of granule size and number (per cell), images were analysed with Fiji (Image J). Micrographs were converted to 8-bit grayscale images and granules were masked by intensity thresholding (Image à Adjust à Threshold). Thresholding was performed manually and was carefully adjusted. Granules were analysed with the *Analyze Particles* tool (Analyzeà Analyse Particles), which selected granules with a size between 0.1 – 10 µm^2^, hence excluding background noise and large aggresomes/aggregates. Granules were approximated as a sphere and area (µm^2^) was converted to average diameter (µm) accordingly. 25-30 cells were analysed per biological replicate. Biological replicates are those which experiments were completed on different days and different passage numbers. SuperPlots were used to convey both cell-level and experimental variability where appropriate. In this case, cell-level values were pooled for each biological replicate, and the mean calculated for each pool. These means were used to calculate the average and SEM bars. Only cells with at least 10 granules were subjected to quantification. Co-localisation analyses were conducted with the DiAna (Distance Analysis) plugin of ImageJ (Fiji).

## Acknowledgements

We acknowledge the use of the SBMS Imaging Facility at the University of Queensland (UQ) in providing the necessary imaging capabilities to complete this project, and the SCMB Mass Spectrometry Facility (UQ), for metabolomics and mass spectrometry. We also acknowledge and thank Milot Mirdita, Sergey Ovhinnokov, Martin Steinegger and the ColabFold team for making their AlphaFold2 modelling pipeline available for public use.

## Author Contributions

**M.J.M:** Conceptualisation; investigation; writing – original draft; writing – review and editing. **Y.Y.Y:** Investigation. **C.C:** Investigation. **J.K.P:** Resources; writing – review and editing. **S.S.M:** Resources; writing – review and editing; **D.C.H.N:** Conceptualisation; resources; supervision; funding acquisition; writing – review and editing.

## Funding

M.J.M is supported by an Australian Government Research Training Program (RTP) scholarship and a Tour de Cure PhD Support Scholarship.

## Conflicts of interest

The authors declare no competing interests.

## Notes

### Competing Interest Statement

The authors have declared no competing interest.

