## Supplementary Figures and Legends for "The microcephaly protein WDR62 regulates cellular purine metabolism through the HSP70/HSP90 chaperone machinery"

**Supplementary Figure Legends**

***Figure S1.*** **AlphaFold-multimer complex predictions for WDR62(FL):BAG2 and WDR62(3936dupC):BAG2. (A)** Complex prediction for WDR62(FL):BAG2, showing interaction between BAG2 with the WD repeat domains and dimerisation domain of WDR62. **(B)** Structure prediction for WDR62(3936dupC):BAG2 which exhibits a truncated C-terminus. **(C)** Boxplot of interphase surface area (Å$^{2}$) calculated by PISA for predicted complexes in **(A)** and **(B).**

***Figure S2.*** **Co-clustering of WDR62, HSP co-chaperones, and DNPB enzymes in response to hyperosmotic stress is not dependent on protein overexpression.** Confocal micrographs of untreated (control) AD293 cells (top row in each set of panels) compared to cells treated with 0.5 M sorbitol for 1 h. Cells transiently transfected with **(A)** WDR62-mCherry and EGFP only or **(B-D)** with mCherry only and **(B)** FGAMS-EGFP, **(C)** PPAT-EGFP or **(D)** GART-EGFP. Fluorescence intensity plots to the right of each set of images demonstrates minimal co-localisation with WDR62 and EGFP, or mCherry and DNPB enzymes. Y-axis represents fluorescence intensity (a.u.), x-axis represents the length of the white line drawn on the ROI (µm). Scale bars on bottom right of merge image represent 20 µm.

***Figure S3.*** **The WDR62 paralog MAPKBP1 does not undergo phase separation or co-localise with DNPB enzymes.** Confocal micrographs of untreated (control) AD293 cells (top row in each set of panels) compared to cells treated with 0.5 M sorbitol for 1 h. Cells transiently transfected with **(A)** MAPKBP1-mCherry only or **(B-D)** with MAPKBP1-mCherry and **(B)** FGAMS-EGFP, **(C)** PPAT-EGFP and **(D)** GART-EGFP. Fluorescence intensity plots to the right of each set of images demonstrates minimal co-localisation with MAPKBP1-mCherry and DNPB enzymes. Y-axis represents fluorescence intensity (a.u.), x-axis represents the length of the white line drawn on the ROI (µm). Scale bars on bottom right of merge image represent 20 µm.

***Figure S4.*** **The condensation of WDR62 requires hyperosmotic stress and unknown bioactive properties of sorbitol. (A)** AD293 cells transfected with WDR62-mCherry and EGFP-G3BP1 and treated with 0.5 M sorbitol or 0.25 M NaCl (both +500 mOsm) for 1 h. Representative confocal micrographs also demonstrate minimal co-localisation between WDR62-mCherry and G3BP-EGFP signal, as indicated by fluorescence intensity plots (*y*-axis represents fluorescence intensity (a.u.), *x*-axis (µm) represents length of white line drawn on ROI image. **(B-C)** WDR62 granule assembly following pre-treatment with the osmo-protectant betaine (20 mM, 1 h). **(E)** AD293 cells expressing WDR62-mCherry treated with millimolar concentrations (5 or 50 mM) sorbitol for 1 h, with or without 0.25 M NaCl. **(F)** Bar graph of proportion (%) of cells with granules with treatments in (A, D-E). Data are the mean of $n \geq 3$

biological replicates. One-way ANOVA with Tukey's multiple comparisons (*p<0.05, **p<0.01, ****p<0.0001, n.s. is p>0.05). All scale bars represent 20 µm. **(G)** Schematic depicting specific findings of experiment. While millimolar concentrations of sorbitol have no effect on granule assembly, combination with 0.25 M NaCl, which alone does not influence WDR62 condensation, induces granule assembly. Osmo-protectants such as betaine abolish WDR62 condensation, suggesting an effect dependent on cell volume changes due to hyperosmotic stress.

***Figure S5.*** **WDR62 granules are associated with mitochondria and microtubules.** Representative confocal micrographs of control (untreated) AD293 cells (top row in each set of panels) compared to cells treated with 0.5 M sorbitol for 1 hour (bottom row in each set). Cells transiently transfected with WDR62-mCherry and stained for **(A)** endogenous alpha-tubulin (α-tubulin) or **(B)** endogenous cytochrome C (Cyt C). Micrographs and white arrows on ROIs demonstrate a clear association between WDR62 granules and both microtubules and mitochondria. Bar graphs on right of each set depict the average percentage of WDR62 granules (mean ± SEM) which overlap with endogenous alpha-tubulin or cytochrome C in individual cells. Pearson's correlation coefficient compared to randomised analysis is also shown. Scale bars on bottom right of merge image represent 20 µm.

***Figure S6.*** **The disordered C-terminus and dimerisation domain of WDR62 are required for granule assembly. (A)** Intrinsic disorder predictions for WDR62. Higher scores indicate greater disorder. **(B)** Schematic depicting full-length WDR62 (WDR62(FL)) (1-1523), and truncated mutants WDR62(N) (1-841), WDR62(C) (842-1523), WDR62(3936dupC) (1-1329), WDR62(842-1329) and WDR62(1290-1523). **(C-H)** Representative images of WDR62 granule under control and sorbitol-treated (0.5 M, 1 h) conditions amongst **(C)** WDR62(FL), **(D)** WDR62(N), **(E)** WDR62(C), **(F)** WDR62(1-1329), **(G)** WDR62(842-1329) and **(H)** WDR62(1290-1523). **(I)** Quantification of proportion (%) of cells from **(C-H)** containing WDR62 granules following sorbitol treatment. Data represent $n$ ≥ 3 biological replicates. One-way ANOVA with Tukey's multiple comparisons. (**** = p<0.0001).

***Figure S7.*** **Loss of WDR62 does not affect purinosome assembly. (A)** Western blot confirming deletion of WDR62 in WDR62 KO cells. **(B)** Confocal micrographs of WT and WDR62 KO AD293 cells expressing FGAMS-mCherry and PPAT-EGFP, before and after sorbitol (0.5 M, 1 h) treatment. Graphs on right of images represent quantifications of no. of granules per cell, proportion of cells with granules, granule diameter (µm) and circularity of purinosomes induced by sorbitol treatment in WT and WDR62 KO cells. Two-tailed unpaired t-test (n.s. is not significant). **(C)** Confocal micrographs of WT and WDR62 KO AD293 cells expressing FGAMS-mCherry and PPAT-EGFP cultured in purine-depleted media for 7 days to induce purinosome assembly.

***Figure S8.*** **WDR62 does not affect the localisation of purinosomes to mitochondria or microtubules.** Confocal micrographs of WT or WDR62 KO AD293 cells transfected with FGAMS-mCherry and stained for either **(A)** endogenous alpha-tubulin or **(B)** endogenous cytochrome C, a mitochondrial marker. Treatment of cells with sorbitol induces the assembly of purinosomes which localise to microtubules and mitochondria. The proportion of mitochondria or microtubule-associated purinosomes does not change following the loss of WDR62.

***Figure S9.*** **WDR62 regulates BAG2 levels. (A)** WT AD293 cells transfected with non-targeting or *WDR62* siRNA expression and immunoblot for BAG2 and HPRT. **(B)** WT AD293 cells transfected with EGFP only or WDR62-EGFP and immunoblot for BAG2 and HPRT. **(C)** WT and WDR62 KO AD293 cells and immunoblot for HSP70, HSP90, STIP1, and DNAJC7.

***Movie S1.*** **WDR62 granules undergo fission and fusion events.**

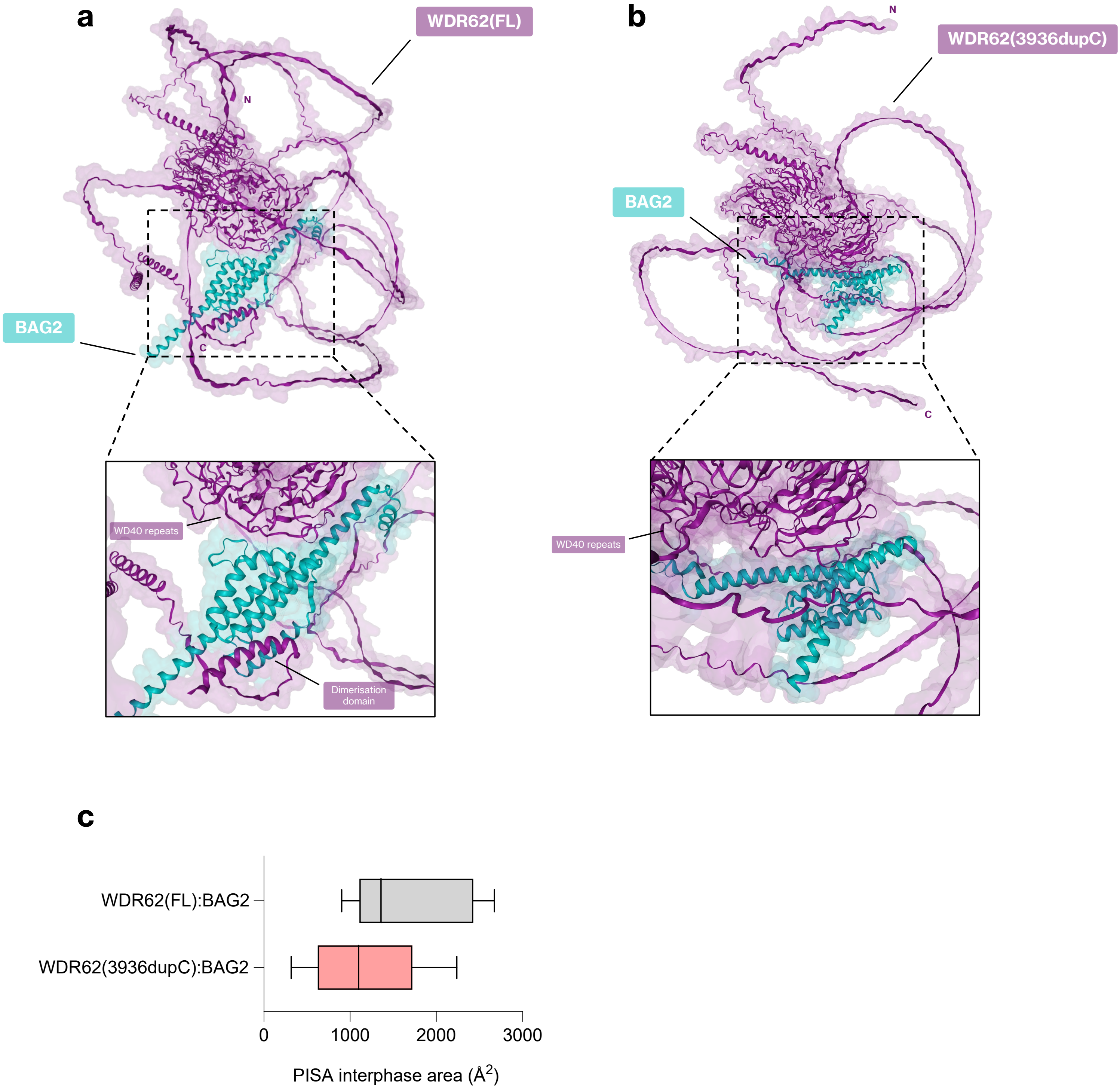
a
WDR62(FL)
N
BAG2
C
WD40 repeats
Dimerisation domain
b
WDR62(3936dupC)
N
BAG2
C
WD40 repeats
c
WDR62(FL):BAG2
WDR62(3936dupC):BAG2
0
1000
2000
3000
PISA interphase area (Å2)

Figure S1

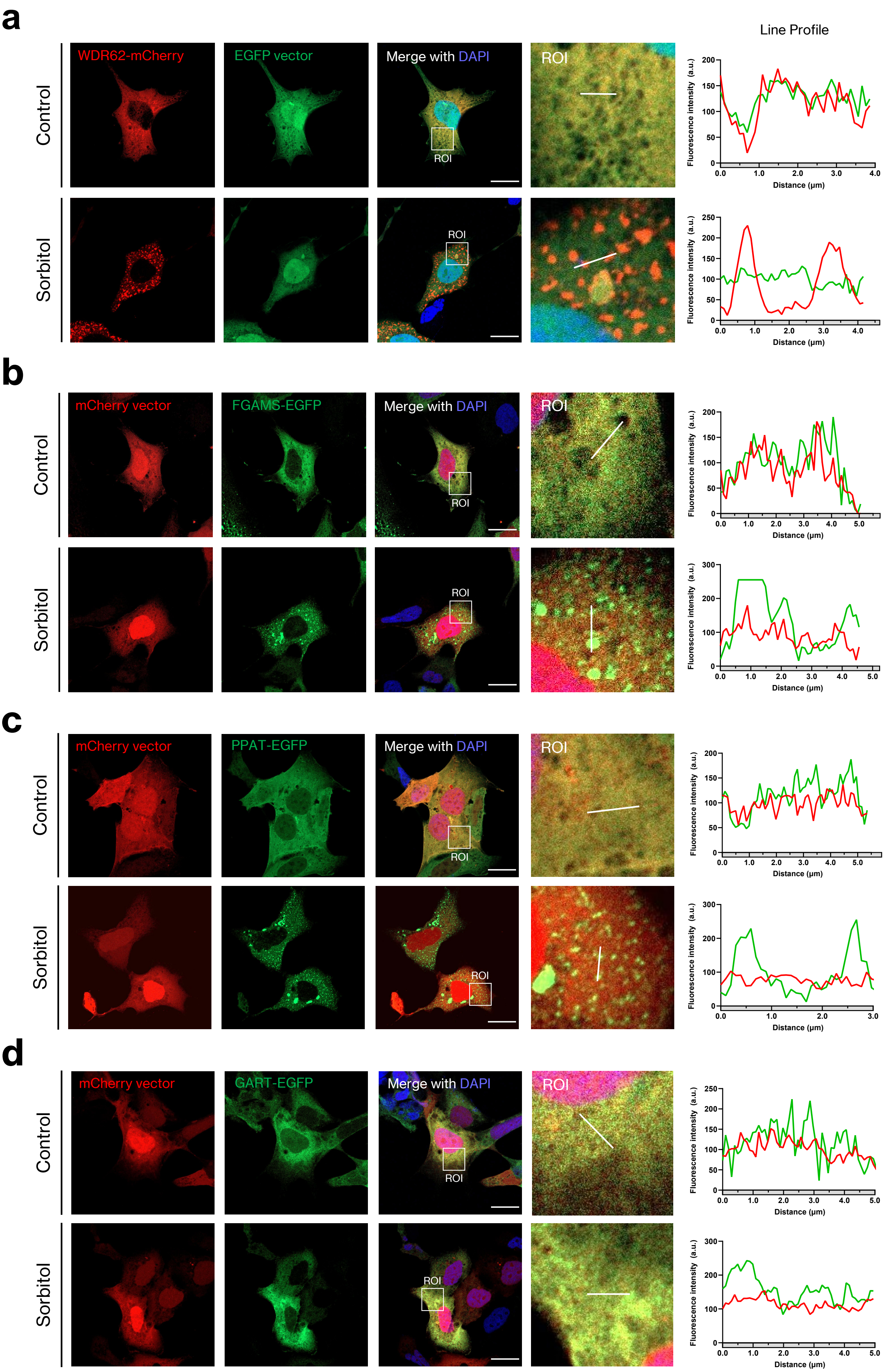
a
Line Profile
WDR62-mCherry
EGFP vector
Merge with DAPI
ROI
Control
Sorbitol
Fluorescence intensity (a.u.)
Distance (μm)
b
mCherry vector
FGAMS-EGFP
Merge with DAPI
ROI
Control
Sorbitol
c
mCherry vector
PPAT-EGFP
Merge with DAPI
ROI
Control
Sorbitol
d
mCherry vector
GART-EGFP
Merge with DAPI
ROI
Control
Sorbitol

Figure S2

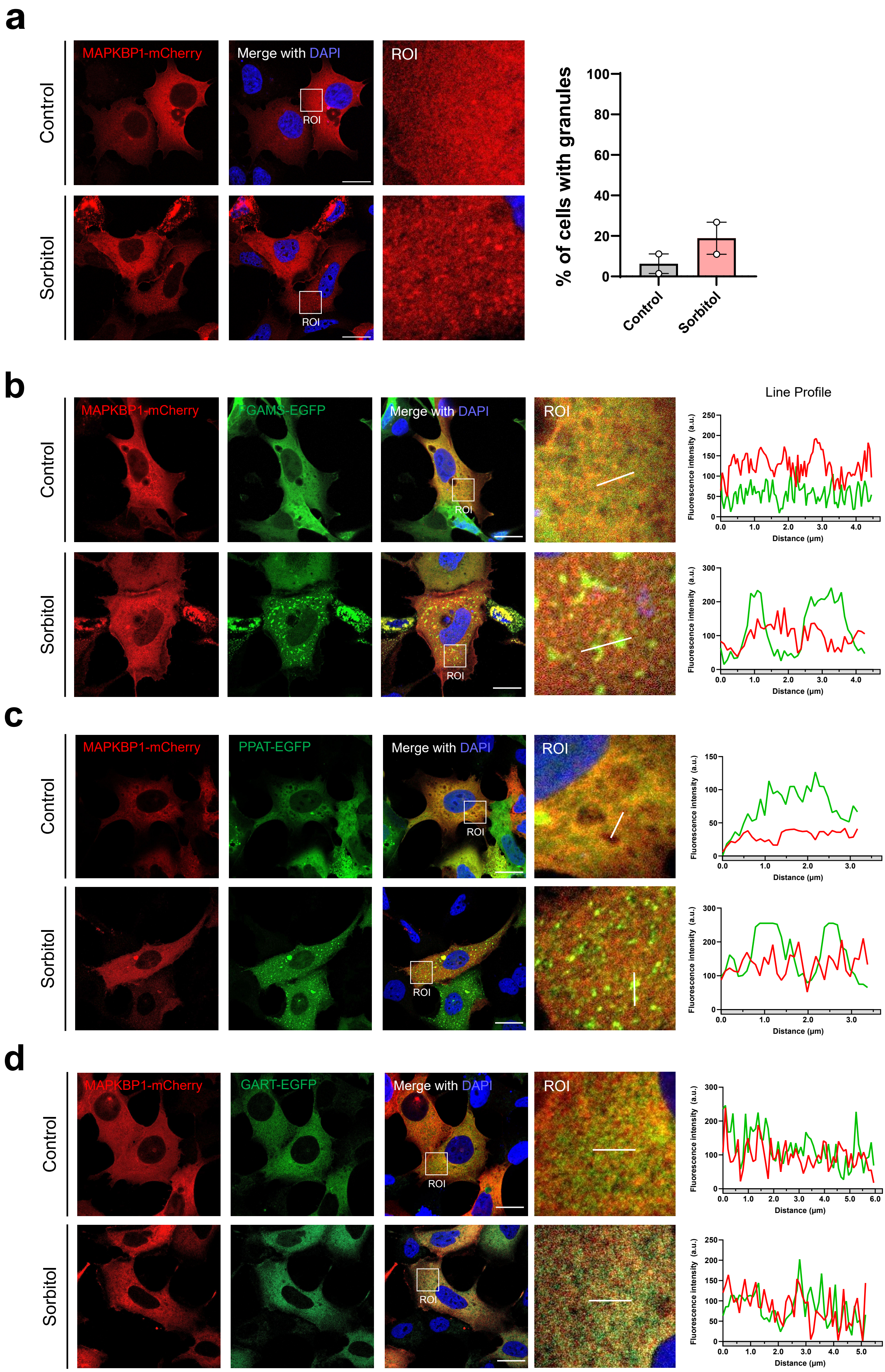

a
MAPKBP1-mCherry
Merge with DAPI
ROI
Control
Sorbitol
% of cells with granules
100
80
60
40
20
0
Control
Sorbitol
b
GAMS-EGFP
Line Profile
Fluorescence intensity (a.u.)
Distance (μm)
c
PPAT-EGFP
d
GART-EGFP


Figure S3

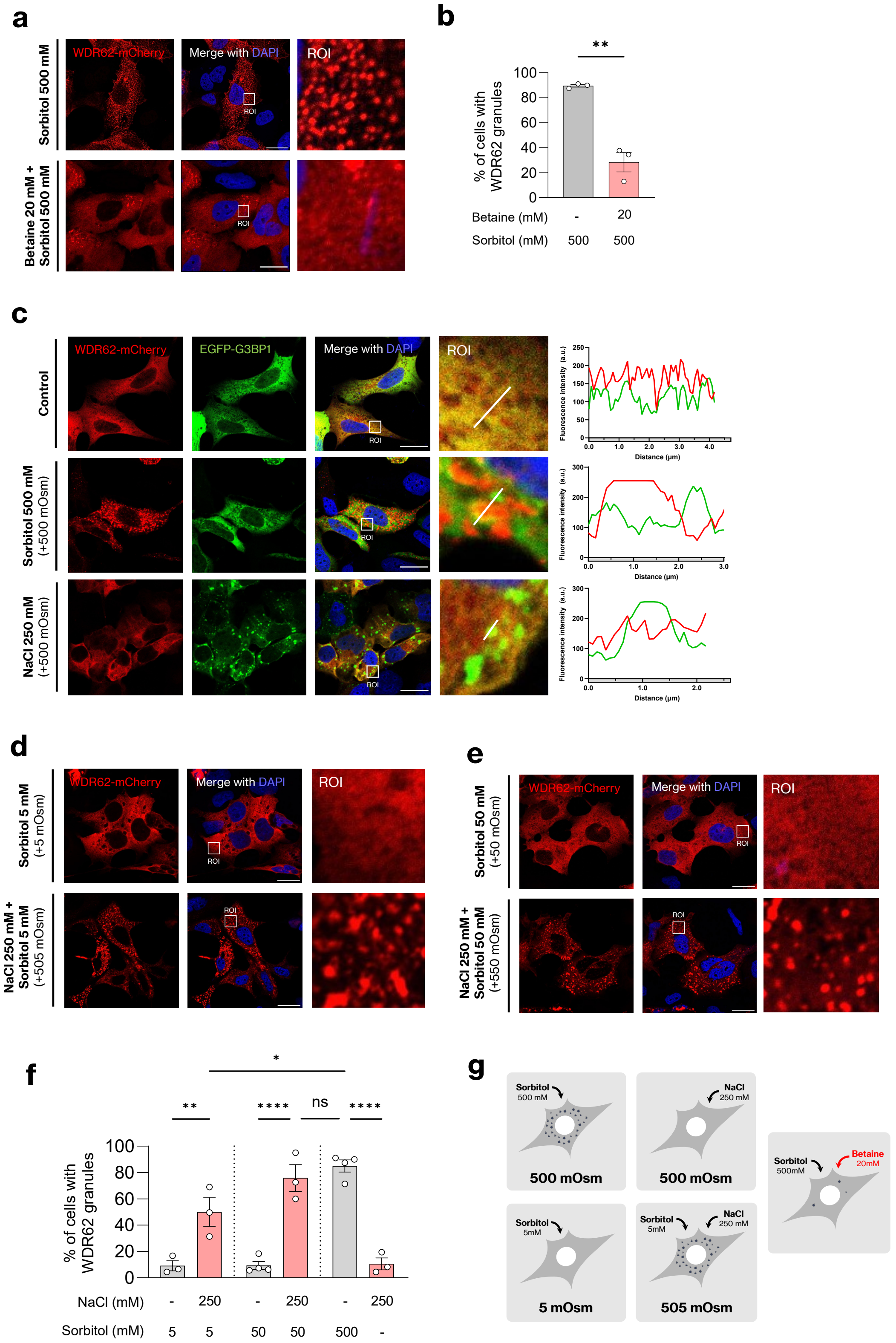
a
Sorbitol 500 mM
Betaine 20 mM + Sorbitol 500 mM
WDR62-mCherry
Merge with DAPI
ROI
b
% of cells with WDR62 granules
Betaine (mM) - 20
Sorbitol (mM) 500 500
c
Control
Sorbitol 500 mM (+500 mOsm)
NaCl 250 mM (+500 mOsm)
WDR62-mCherry
EGFP-G3BP1
Merge with DAPI
ROI
Fluorescence intensity (a.u.)
Distance (μm)
d
Sorbitol 5 mM (+5 mOsm)
NaCl 250 mM + Sorbitol 5 mM (+505 mOsm)
e
Sorbitol 50 mM (+50 mOsm)
NaCl 250 mM + Sorbitol 50 mM (+550 mOsm)
f
% of cells with WDR62 granules
NaCl (mM) - 250 - 250 - 250
Sorbitol (mM) 5 5 50 50 500 -
ns
g
Sorbitol 500 mM
500 mOsm
NaCl 250 mM
500 mOsm
Sorbitol 5mM
5 mOsm
Sorbitol 5mM
NaCl 250 mM
505 mOsm
Sorbitol 500mM
Betaine 20mM

Figure S4

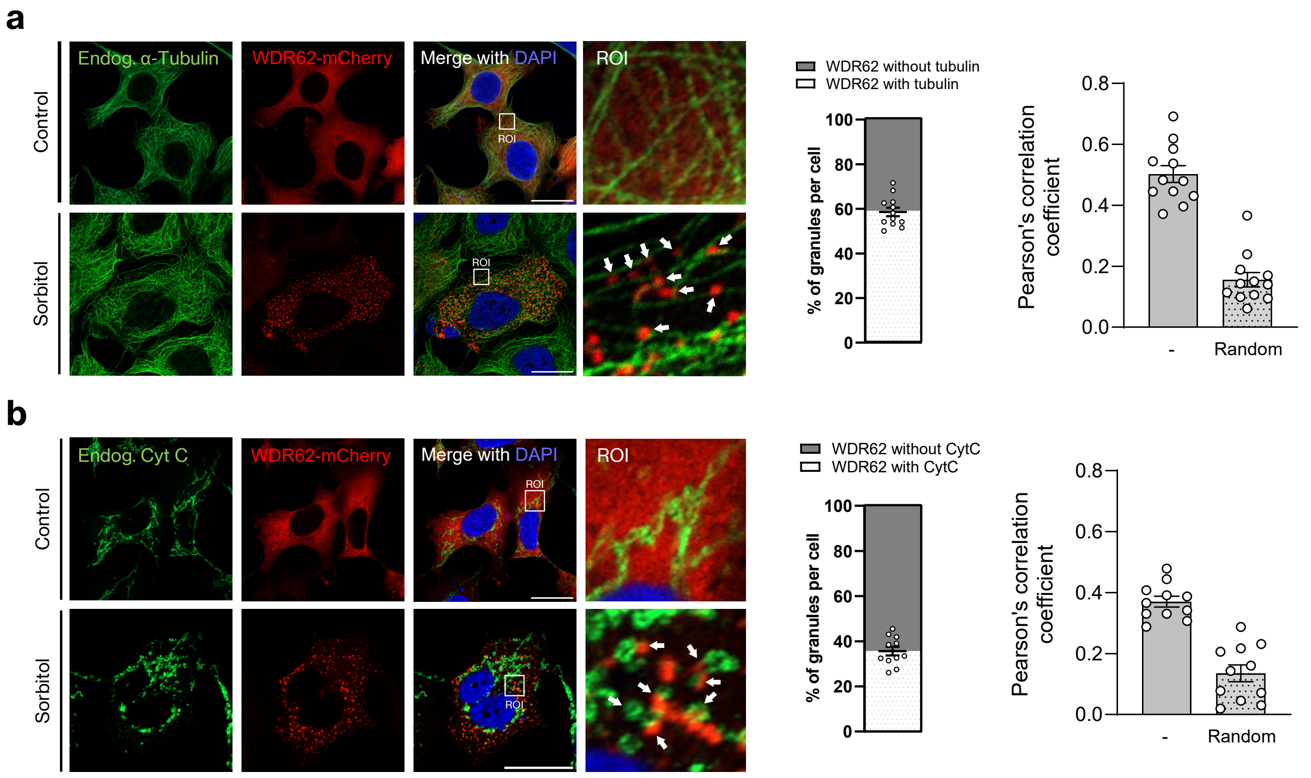
a
Endog. α-Tubulin
WDR62-mCherry
Merge with DAPI
ROI
Control
Sorbitol
WDR62 without tubulin
WDR62 with tubulin
% of granules per cell
Pearson's correlation coefficient
-
Random
b
Endog. Cyt C
WDR62-mCherry
Merge with DAPI
ROI
Control
Sorbitol
WDR62 without CytC
WDR62 with CytC
% of granules per cell
Pearson's correlation coefficient
-
Random

Figure S5

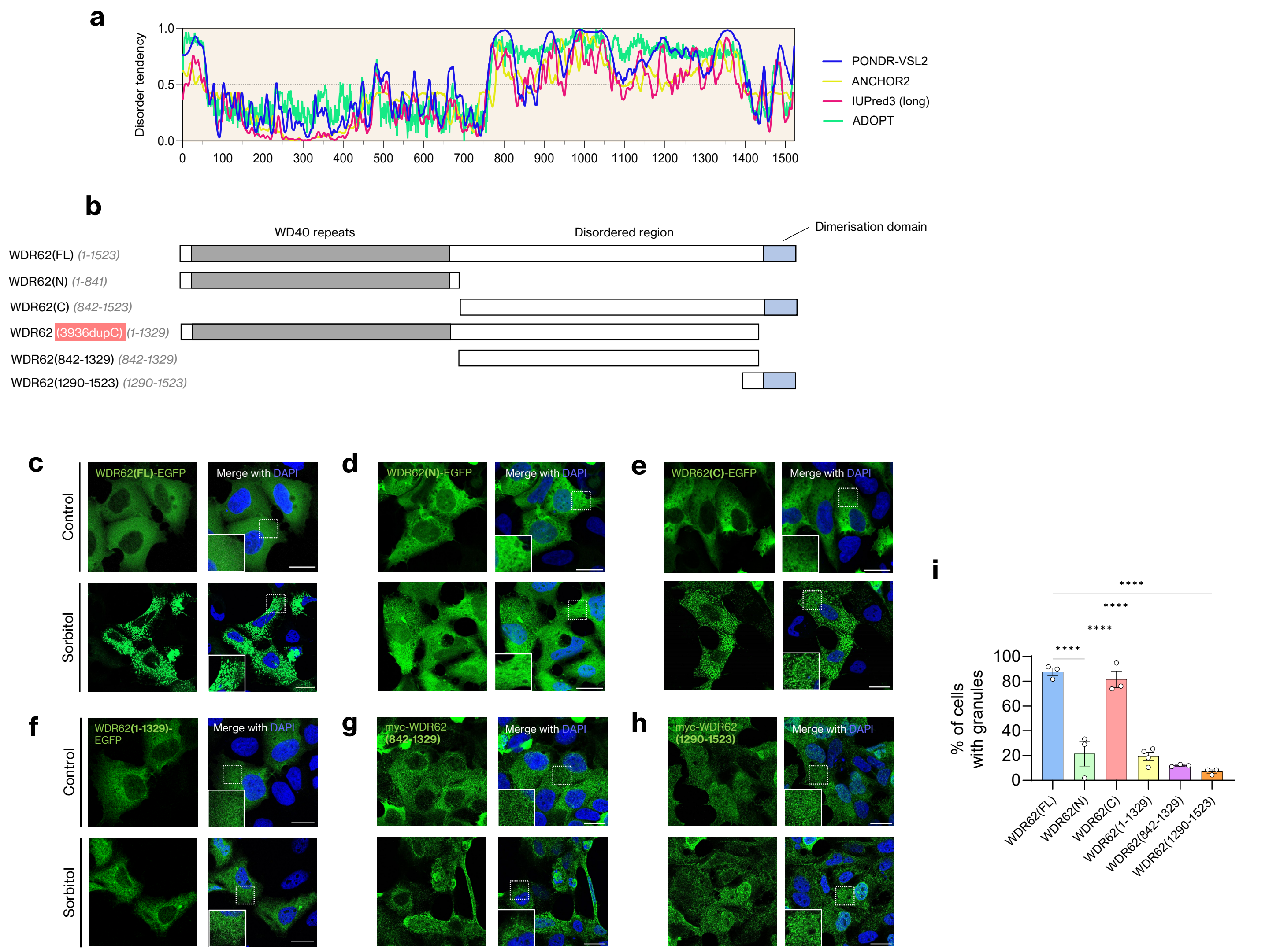
a
Disorder tendency
1.0
0.5
0.0
0
100
200
300
400
500
600
700
800
900
1000
1100
1200
1300
1400
1500
PONDR-VSL2
ANCHOR2
IUPred3 (long)
ADOPT
b
WD40 repeats
Disordered region
Dimerisation domain
WDR62(FL) (1-1523)
WDR62(N) (1-841)
WDR62(C) (842-1523)
WDR62 (3936dupC) (1-1329)
WDR62(842-1329) (842-1329)
WDR62(1290-1523) (1290-1523)
c
d
e
f
g
h
Control
Sorbitol
WDR62(FL)-EGFP
WDR62(N)-EGFP
WDR62(C)-EGFP
WDR62(1-1329)-EGFP
myc-WDR62 (842-1329)
myc-WDR62 (1290-1523)
Merge with DAPI
i
****
% of cells with granules
100
80
60
40
20
0
WDR62(FL)
WDR62(N)
WDR62(C)
WDR62(1-1329)
WDR62(842-1329)
WDR62(1290-1523)

Figure S6

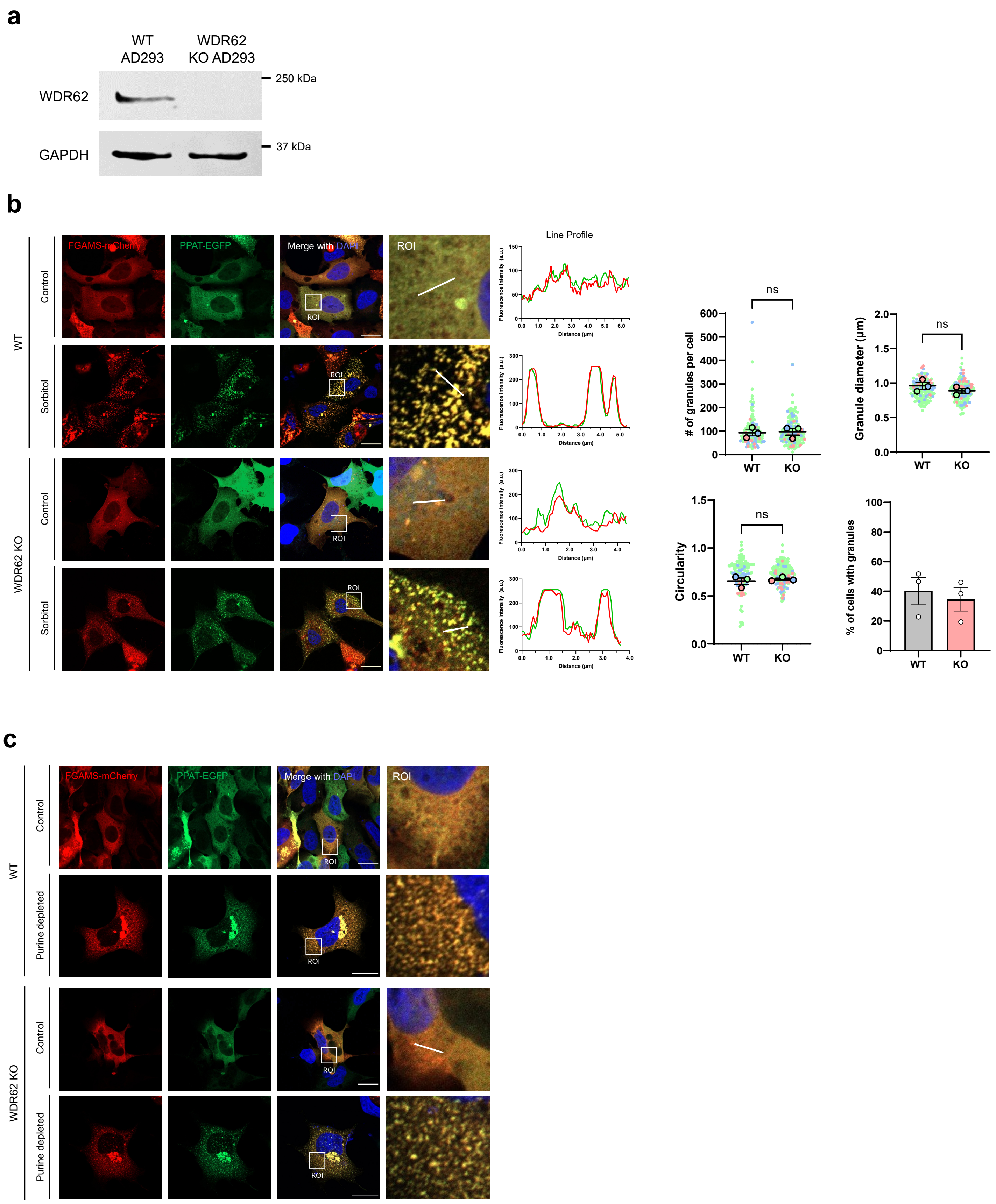
a
WT AD293
WDR62 KO AD293
WDR62
GAPDH
250 kDa
37 kDa
b
WT
WDR62 KO
Control
Sorbitol
FGAMS-mCherry
PPAT-EGFP
Merge with DAPI
ROI
Line Profile
Fluorescence intensity (a.u.)
Distance (µm)
### of granules per cell
Granule diameter (µm)
Circularity
% of cells with granules
ns
KO
c
Purine depleted

Figure S7

**a**

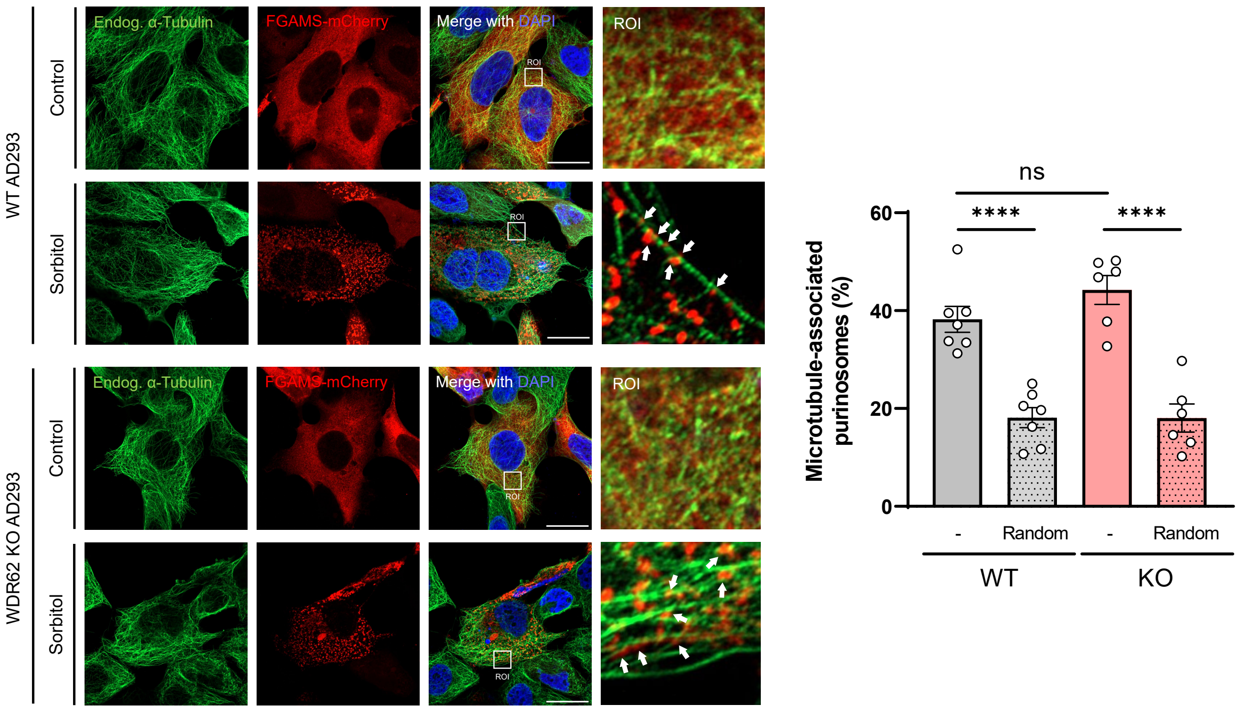
WT AD293
WDR62 KO AD293
Control
Sorbitol
Endog. α-Tubulin
FGAMS-mCherry
Merge with DAPI
ROI
Microtubule-associated purinosomes (%)
0
20
40
60
ns
****
****
-
Random
WT
KO

**b**

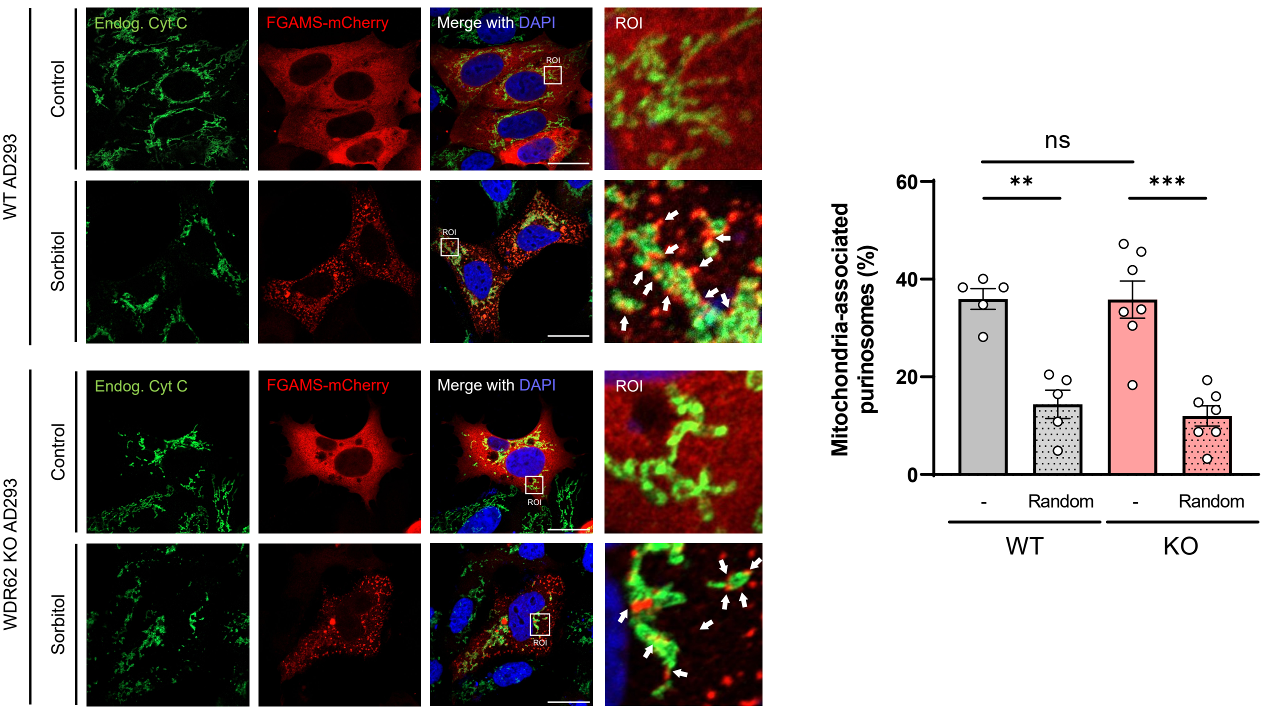
WT AD293
WDR62 KO AD293
Control
Sorbitol
Endog. Cyt C
FGAMS-mCherry
Merge with DAPI
ROI
Mitochondria-associated purinosomes (%)
0
20
40
60
ns
**
***
-
Random
WT
KO

Figure S8

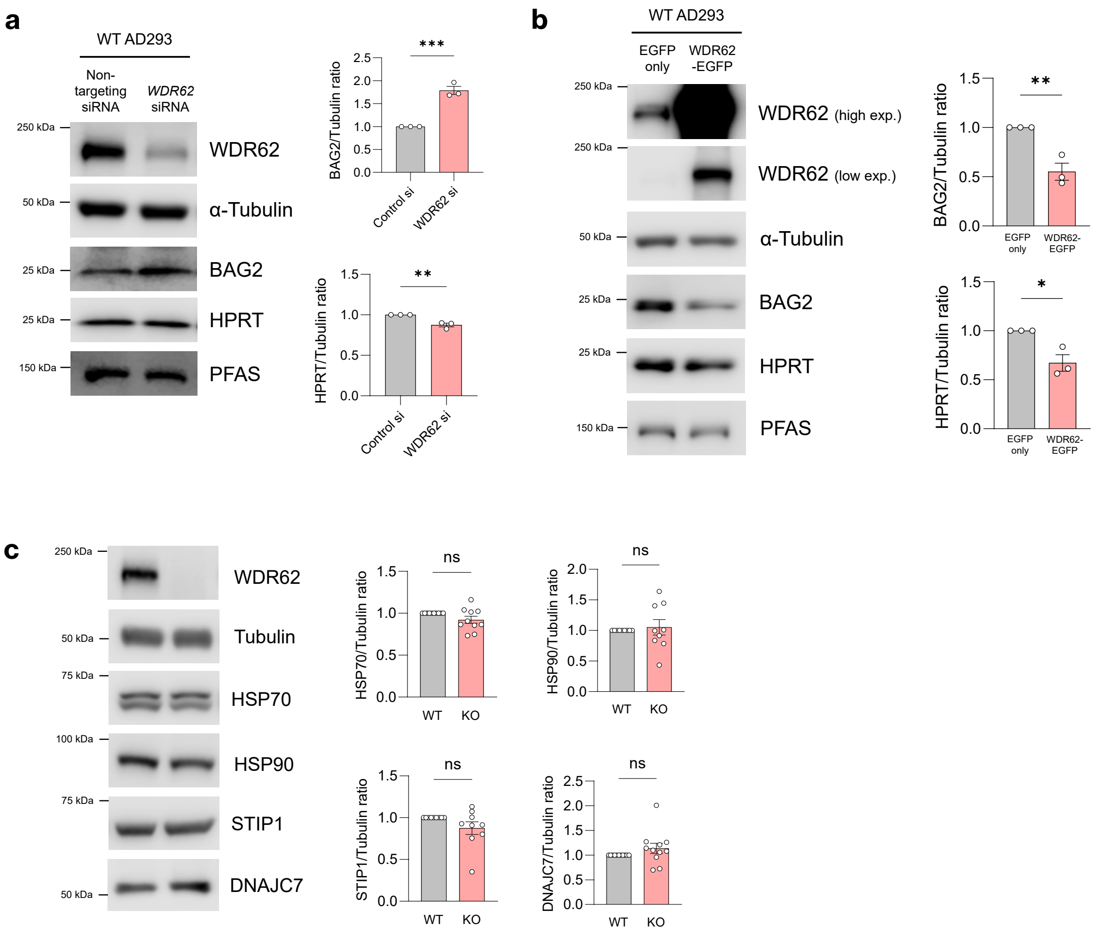
a
WT AD293
Non-targeting siRNA
WDR62 siRNA
250 kDa
WDR62
50 kDa
α-Tubulin
25 kDa
BAG2
25 kDa
HPRT
150 kDa
PFAS
BAG2/Tubulin ratio
***
Control si
WDR62 si
HPRT/Tubulin ratio
**
b
WT AD293
EGFP only
WDR62 -EGFP
WDR62 (high exp.)
WDR62 (low exp.)
α-Tubulin
BAG2
HPRT
PFAS
BAG2/Tubulin ratio
**
HPRT/Tubulin ratio
*
c
WDR62
Tubulin
75 kDa
HSP70
100 kDa
HSP90
STIP1
DNAJC7
HSP70/Tubulin ratio
HSP90/Tubulin ratio
STIP1/Tubulin ratio
DNAJC7/Tubulin ratio
ns
WT
KO

Figure S9
